# RatDISCO, a tissue clearing and immunolabelling protocol for large rat brains

**DOI:** 10.1101/2025.08.17.670723

**Authors:** Kirsty J. Craigie, Sally Till, Britt van de Gevel, Brianna Vandrey, Patrick Strangward, Dirk Sieger, Alfredo Gonzalez-Sulser, Peter Kind, Nathalie Rochefort, Ian Duguid, Cristina Martinez-Gonzalez

## Abstract

RatDISCO is a simple, cost-effective, and reproducible tissue-clearing protocol optimised for immunolabeling in adult rat brains. It enables robust detection of diverse neuronal subtypes, glial populations, and vasculature, overcoming key limitations in antibody penetration and optical transparency. It is compatible with virally labelled-and transgenic mice, as well as human-derived brain organoids, highlighting its versatility across species and models. Functionally, it enables the detection of activity-dependent markers after behaviour, verified in a rat model of neurodevelopmental disorders, the Fragile X syndrome. These findings highlight RatDISCO as a broadly applicable tool for investigating neuroanatomical and functional alterations in rat models of disorders.

**Highlights:**

- Optimised for immunolabeling in adult rat brains
- Enables detection of neurons, glial cells, and vasculature
- Achieves high tissue transparency without requiring specialised equipment.
- Compatible with virally labelled mice and human brain organoids
- Facilitates mapping of activity-dependent markers in rat models of neurodevelopmental disorders

## Introduction

Unbiased three-dimensional (3D) mapping of the neurochemical architecture is essential for understanding brain function in both health and disease^1^. Rodent models, especially mice, have been central to this effort, largely due to the availability of genetic tools that enable precise manipulation of gene expression and function. However, rats offer several advantages, including larger brains, more complex behavioural repertoires, and increasing access to genetically modified strains^2,3^. These attributes make rats particularly valuable for studying genetically linked conditions, including developmental disorders^4–9^.

Traditional histological techniques rely on slicing tissue into thin sections for immunolabeling, which can introduce artifacts and lead to tissue loss, compromising 3D structural integrity. Tissue-clearing methods have advanced considerably, improving optical transparency and reducing light scattering in intact samples. Protocols such as 3DISCO^10^, CLARITY^11,12^, 2,2’-thiodiethanol-TDE^13^, and ethyl cinnamate-ECi^14^ have enabled whole-organ imaging, yet consistent antibody penetration in large, intact tissues remains a major challenge^15^. Techniques like iDISCO+^16^, SeeDB^17^, and SHANEL^18^ improve antibody penetration but are primarily optimised for mouse or human tissue.

Despite the increasing importance of rat models, no standardised method currently exists for immunolabeling and clearing adult rat brains. To address this gap, we developed RatDISCO, a robust, solvent-based clearing method that incorporates antigen retrieval steps to enhance antibody penetration in adult rat tissue. We evaluated RatDISCO’s performance in tissue transparency, shrinkage, and immunolabeling efficacy, and benchmarked it against established protocols. We demonstrated its utility for labelling neuronal subtypes, glial cells, and vasculature in adult rat brains using immunohistochemistry, and tested its compatibility with other species, including mice and human-derived brain organoids. Finally, we used RatDISCO to evaluate cFOS-positive neurons in the amygdala, used as a proxy for experience-dependent neural activity^19–21^, in a rat model of Fragile X syndrome^22^, demonstrating its utility for functional brain mapping in genetically linked disorders.

## Results

### RatDISCO achieves optical transparency, immunolabelling and isotropic shrinkage in adult rat brains

RatDISCO is a hydrophobic tissue clearing method^1,23–25^ that consists of 4 steps to achieve immunolabelling and optical clearing of adult rat brains. It combines antigen retrieval to expose antigens masked by aldehyde fixation-crosslinking before immunolabelling^26^ (Figure 1A). To further enhance this, we use methanol^27,28^ and surfactants^29,30^ to later allow the penetration of the antibodies through the large volume of the rat brain (Figure 1B). We then dehydrate and delipidate the tissue using increasing concentrations of tetrahydrofuran (THF)^10,31,32^ and dichloromethane (DCM)^33,34^. Finally, we use dibenzyl ether (DBE) for refractive index (RI) matching^10,31,32^ and ethyl cinnamate-ECi^14^ for tissue storage and light-sheet microscopy imaging.

**Figure 1.**
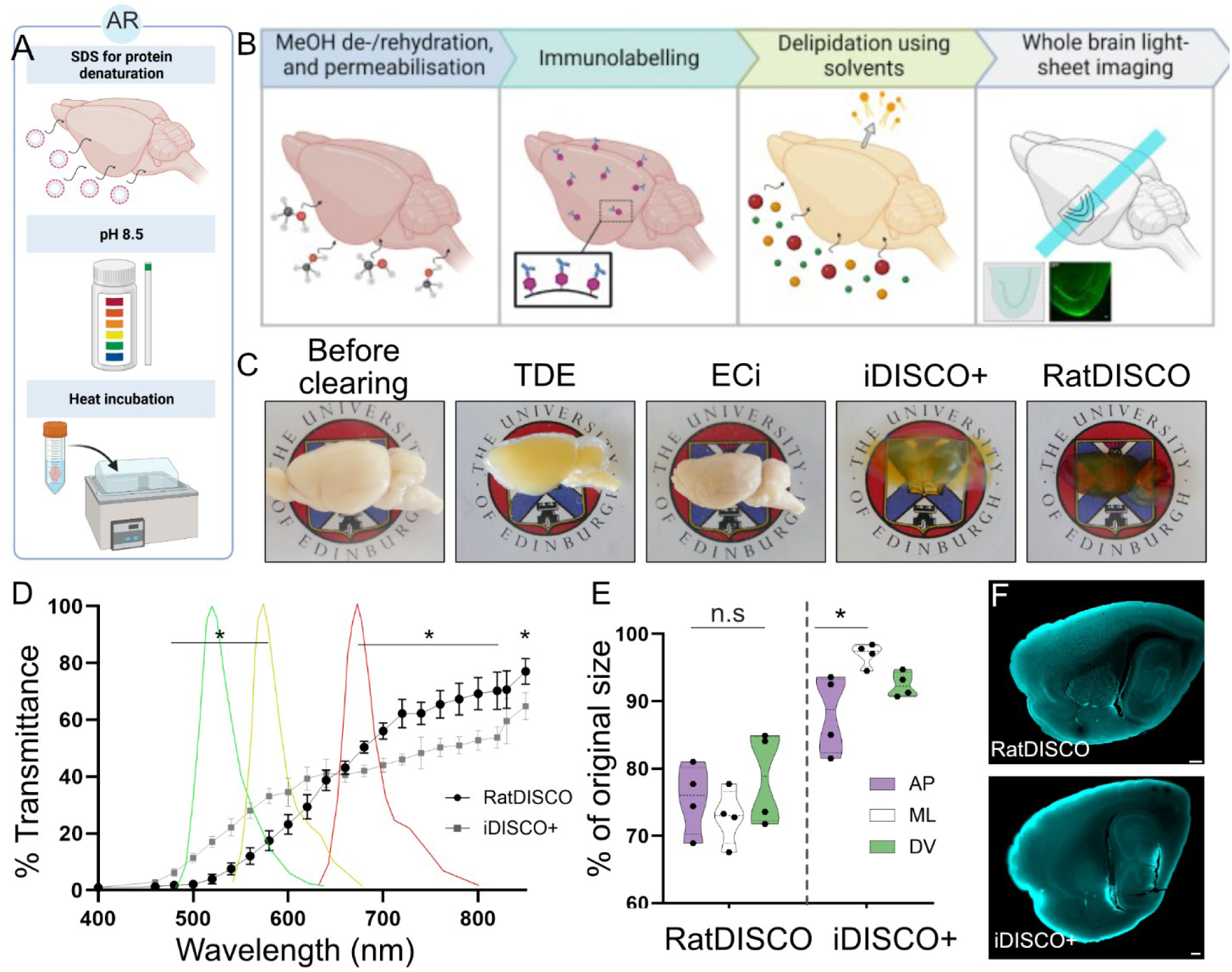
RatDISCO immunolabelling and optical clearing pipeline. Schematic drawing of the RatDISCO protocol. A. Antigen retrieval (AR?) incubation step: SDS at pH 8.5 and 80°C (optional step 1). B. RatDISCO steps: methanol (MeOH) de/rehydrations, permeabilisation and blocking (1), immunolabelling (2), delipidation and optical clearing (3), and imaging using light sheet microscopy (4). C. Images of rat brain hemispheres before clearing and optically cleared using TDE and ECi (paired), and iDISCO+ and RatDISCO (paired). D. Comparison of the percentage of light transmittance in tissue cleared with RatDISCO (black circles; mean and SD) or iDISCO+ (grey squares; mean and SD). Transmittance measurements between RatDISCO and iDISCO+ at specific wavelengths were compared using a two-tailed paired t-test (P < 0.05*). Means, SD, P values and confidence levels per measured wavelength are described in Table 1. Green, yellow and red lines represent the emission spectra of Alexa dyes 488, 546 and 647, respectively. D. Isotropic vs anisotropic tissue shrinkage was assessed by measuring dimensional changes along the anterior-posterior (AP-purple), mediolateral (ML-white) and dorsoventral (DV-green) axes (shown as truncated violin plots; median and quartiles: dashed lines; original tissue size per represented as 100%). RatDISCO-cleared tissue shrank isotropically with no significant differences (n.s) in shrinkage across anatomical axes (ANOVA: F (2,9) = 1.095; P = 0.38). iDISCO+cleared tissue showed anisotropic shrinkage with significant differences between the AP and ML axes (ANOVA: F (2,9) = 5.731; P < 0.02). Tukey’s multiple comparison test values are reported in Table 2. *P < 0.05. F. Light-sheet microscope images of the globus pallidus from paired hemispheres immunolabelled against FoxP2 (cyan) and cleared with RatDISCO (above) and iDISCO+ (below). Scale bars D above: 250 µm; below 500 µm.

We compared the tissue transparency of adult rat brain paired hemispheres cleared using RatDISCO to three existing clearing methods standardised in mice (Figure 1C). We used iDISCO+ as it has been reported to offer enhanced immunolabelling capabilities in the whole mouse brain to that of its predecessor iDISCO. We also compared TDE and ECi clearing, as they both have been reported as effective clearing methods in mice and offer greater biological safety in comparison to THF and DCM solvents^3,4,9,18^.

Both RatDISCO and iDISCO+ rendered tissue optically transparent, but with distinct physical and visual characteristics (Figure 1C). RatDISCO-cleared tissue had an amber-like tint, and formed a resin-like solid that was easy to handle and image. In contrast, iDISCO+ cleared tissue appeared lighter in hue and was soft, brittle, and prone to breaking, making it difficult to manage. Surprisingly, TDE-and ECi-cleared tissue did not achieve transparency, even after a prolonged incubation of several months (Figure 1C), and were therefore excluded from further analysis.

To investigate the transparency achieved with RatDISCO and iDISCO+, we measured and compared the percentage of light transmitted through paired brain hemispheres from the same animal, cleared using either method, across wavelengths from 450 to 850 nm at 20-60 nm intervals (n=4 rats; Figure 1D). We observed a higher percentage of light transmittance in iDISCO+ cleared tissue compared to RatDISCO at shorter wavelengths, specifically between 450 and 580 nm. Both methods showed comparable light transmittance at wavelengths between 600 and 660 nm, and at 830 nm. However, at longer wavelengths (680 – 820 nm and 850 nm), RatDISCO-cleared tissue exhibited significantly higher light transmittance than iDISCO+ cleared tissue (Figure 1E; two-tailed paired t-test P <0.05; full statistical details are provided in Supplementary Table 1). These findings indicate that iDISCO+ cleared tissue transmits light more effectively at shorter wavelengths, whereas RatDISCO performs better at longer wavelengths.

Our next question was whether RatDISCO-cleared tissue shrank at the same ratio as other solvent-based optical clearing methods. RatDISCO uses sequential methanol dehydration and solvent incubation steps to permeabilise and delipidate the tissue while preserving protein localisation, resulting in tissue shrinkage. iDISCO+ is reported to produce an overall shrinkage of 11% compared to 50% using the original 3DISCO clearing protocol^10,16^. To compare shrinkage between RatDISCO and iDISCO+, we hemisected 4 rat brains along the sagittal plane and measured the percentage of shrinkage in paired hemispheres cleared with each method. tissue dimensions were measured along the anterior-posterior (AP), mediolateral (ML) and dorsoventral (DV) axes before and after clearing with RatDISCO or iDISCO+ (Figure 1E).

RatDISCO-cleared tissue exhibited an overall shrinkage of 24.3%, resulting in measurements of 75.7% ± 5.5% (mean ± SD) of its original size. We observed no significant differences in shrinkage across the individual AP, ML and DV anatomical axes for RatDISCO-cleared tissue, suggesting isotropic shrinkage of the tissue. In contrast, iDISCO+ cleared tissue showed significantly less overall shrinkage (7.5%), retaining 92.5% ± 5% (mean ± SD) of its original size, but exhibited a significant difference in shrinkage between the AP and ML axes, with no significant change along the DV axis (Figure 1F; RatDISCO ANOVA: F (2,9) = 1.095; P = 0.3753. iDISCO+ ANOVA: F (2,9) = 5.731; P = 0.0248; Tukey multiple comparisons described in Supplementary Table 2).

These results indicate that RatDISCO-cleared tissue undergoes isotropic shrinkage, with no significant differences across the three anatomical axes. In contrast, iDISCO+ cleared tissue exhibits anisotropic shrinkage, with greater reduction along the AP axis compared to the ML axis, while the DV axis remains largely unaffected. This highlights the distinct shrinkage patterns associated with the two clearing methods.

Finally, we asked whether immunolabelling was achievable and comparable between RatDISCO and iDISCO+ (Figure 1F). We used an antibody against the transcription factor Forkhead box protein P2 (FoxP2)^35,36^, due to its brain-wide expression and known compatibility with iDISCO+ in mice^28^. FoxP2 is expressed in neurons in superficial and deeper areas of the rat brain, including the cortex, globus pallidus and the amygdala^20,21^. We detected FoxP2-positive neurons (Figure 1F, cyan) in the expected regions in tissue cleared with RatDISCO, whereas labelling was noticeably reduced in tissue processed with iDISCO+.

### RatDISCO achieves deep antibody labelling in adult rat brains

To quantitatively compare FoxP2 immunolabelling between RatDISCO and iDISCO+ clearing methods, we processed brain hemispheres from the same animals using each clearing method. We acquired sagittal light-sheet microscope images at both low and high magnification (Figures 2 A-B and C-D, respectively). Using commercially available software (Arivis Vision 4D), we created a custom 3D template atlas of the rat amygdala to spatially register and quantify FoxP2-immunolabelled nuclei in the acquired light-sheet data sets (Supplementary Figure 1). We based our atlas on the 2D Paxinos and Watson^37^ rat brain atlas to create a 3D atlas that delineates the four major basolateral amygdala complexes: the basal, basomedial, lateral, and central complexes^38^ (Figure 2E). Given the tissue shrinkage differences between RatDISCO and iDISCO+ clearing methods, we adjusted the size of the atlas accordingly and normalised measurements using cell density (labelled nuclei per volume) to enable meaningful comparisons between methods and across animals.

**Figure 2.**
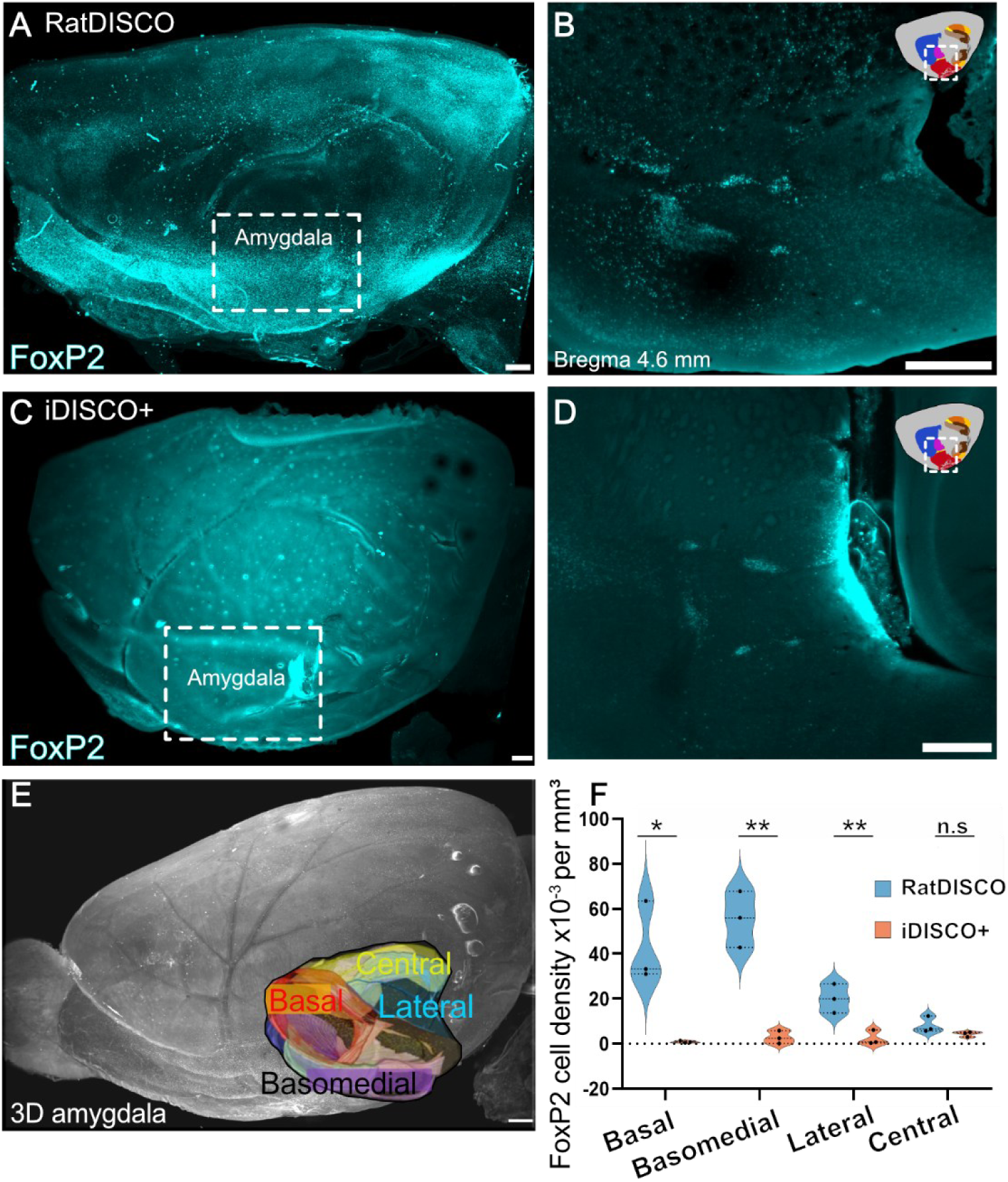
Immunolabelling comparison between RatDISCO and iDISCO+. Light-sheet microscope images of paired rat brain hemispheres immunolabelled against FoxP2 and cleared with RatDISCO (A, B) or iDISCO+ (C, D). A, C. whole-brain maximum intensity projections. B, D. Single optical plane images of the amygdala. E. Schematic of our custom-made 3D template atlas for the rat amygdala delineating its four major basolateral complexes: the basal, basomedial, lateral, and central complex superimposed on the whole-brain. F. Violin plots showing the density of FoxP2-positive neurons in amygdala subregions from brains cleared with RatDISCO (blue) or iDISCO+ (orange). Circle dots represent individual animals; dashed lines show median and quartile values. (Two-tailed t-test; basal: P = 0.02, t=4.0, df=4; basomedial: P = 0.001, t=7.1, df=4; lateral: P = 0.01, t=4.3, df=4, and central amygdala complex: P = 0.14; t=1.8, df=4.). n.s > 0.05; *P < 0.05; **P < 0.01. Scale bars A, B, F: 250 µm; C, D: 500 µm.

To quantify immunolabelled FoxP2-positive nuclei in amygdala subregions, we used a customised “blob finder” analysis pipeline in Arivis Vision 4D. We manually validated parameters to automatically detect and quantify nuclei in 3D light-sheet data sets (Supplementary Figure 2). We applied this pipeline to FoxP2-immunolabelled brain hemispheres from the same animal cleared with RatDISCO or iDISCO+. In RatDISCO cleared tissue, FoxP2-positive cell densities (x10^-3^ per mm^3^) were 43 ± 18 in the basal complex, 56 ± 13 in the basomedial, 8.2 ± 3.6 in the central, and 20 ± 6.4 in the lateral amygdala. In the iDISCO+ cleared tissue, densities were lower: 0.84 ± 0.43 in the basal, 2.8 ± 2.9 in the basolateral, 4.3 ± 1.1 in the central, and 2.4 ± 3.3 cells x10^-3^ per mm^3^ in the lateral amygdala (Figure 2F).

Statistical analysis revealed significant differences in FoxP2-positive cell density between RatDISCO and iDISCO+ in the basal (P = 0.0163, t = 4.0), basomedial (P = 0.0021, t = 7.1), and lateral (P = 0.0131, t = 4.3) complexes of the amygdala, but not in the central complex (P = 0.1445; t = 1.8)(Two-tailed unpaired t-tests; df = 4 for all comparisons). Together, these findings suggest that RatDISCO provides substantial improvements in detection of FoxP2-immunolabelled neurons in rat tissue compared to iDISCO+.

### RatDISCO allows labelling of distinct neuronal populations, microglia and vasculature in the rat brain

To demonstrate the versatility of RatDISCO, we immunolabelled transcription factors expressed by distinct neuronal populations in the rat brain, including cFOS, FoxP2^39,40^ and COUP TF1-interacting protein 2^41^ (CTIP2; Figure 3). cFOS is an immediate early gene rapidly and transiently induced by extracellular stimuli and is widely used as a proxy for neuronal activity in the brain^19,21,42,43^ (Figure 3 A-C). FoxP2 is broadly expressed across the brain, with strong enrichment in the globus pallidus, internal capsule of the amygdala, and the inferior olive^36,40,44–46^ (Figure3 D-F). CTIP2 shows more region-specific expression, particularly in the hippocampus, amygdala and layer 5 of the medial entorhinal cortex^47–49^ (Figure 3 G-I).

**Figure 3.**
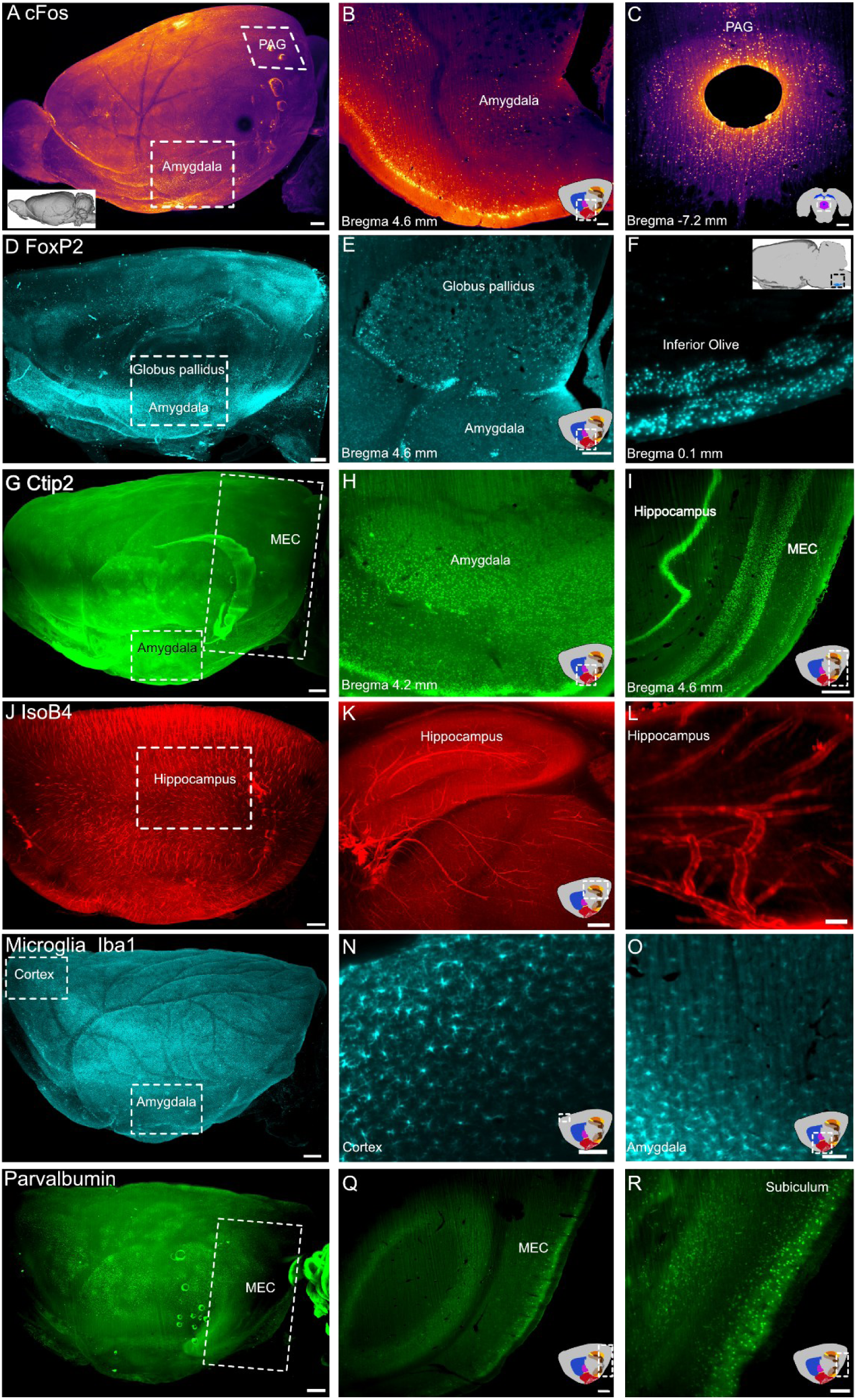
**RatDISCO allows the visualization of several neuronal, glial and blood vessel markers in whole rat brains**. Light-sheet microscopy images of tissue immunolabelled against cFOS (A-C Inferno LUT), FoxP2 (D-F, cyan), Ctip2 (G-I, green), IsoB4 (J-L, red, blood vessels), Iba1 (M-P; microglia, cyan), Parvalbumin (P-R, green). PAG: Periaqueductal gray. Schematic representations of the approximate area and anatomical level on the inset. A, D, G, J, M, P. Maximum intensity projections; scale bars 500 µm. B, C, F, Q, R. Single optical slices; scale bars 100 µm. Scale bars E, H, I, K: 250 µm and L, N, O: 50 µm.

We also used the lectin IsoB4, which binds to endothelial cells^50^ to visualise the detailed vasculature of the rat brain (Figure 3 J-L). To visualise microglia and border macrophages, we immunolabelled against the ionised calcium-binding adaptor molecule 1 (Iba1)^51,52^ (Figure 3 M-O). Fast-spiking GABAergic interneurons were identified by immunolabelling for parvalbumin^53,54^ (Figure 3 P-R). Finally, we also adapted an RNAscope *in situ* hybridization protocol^55^ to evaluate the compatibility of RatDISCO with RNA detection. To test this, we used a probe targeting POLR2A (DNA-directed RNA polymerase II subunit RPB1), a well-established low-copy, yet robust, positive control for detecting low-expressing RNA targets. We detected POLR2A signal throughout the brain, including in the cerebellum, brainstem, cortex and amygdala (Supplementary Figure 3). Together, these findings demonstrate the versatility of RatDISCO for whole-brain imaging in rats, enabling reliable labelling of diverse neuronal subtypes, glial cells, vasculature, and RNA transcripts. Moreover, they underscore the method’s broad applicability for studying complex neuroanatomical and molecular features across multiple brain regions.

### RatDISCO is compatible with mouse brain tissue and organoids

Next, we tested whether RatDISCO could also be applied to mouse brains. We found that is successfully enabled immunolabelling of MECP2, a protein associated with Rett syndrome^56^, as well as cFOS (in an acute intrahippocampal kainate temporal lobe epilepsy model)^57,58^, tyrosine hydroxylase (TH)^59^-and FoxP2-positive across multiple brain regions (Figure 4 A-C and G-H, respectively). RatDISCO also enabled visualisation of virally-labelled axonal projections from the medial and lateral entorhinal cortex (MEC and LEC, respectively) in mice. To label projections from the MEC to the hippocampus, we injected a Cre-dependent adeno-associated virus (AAV) expressing mCherry into the MEC of Sim1^Cre^ transgenic mice to label projections to the hippocampus (Figure 4 D-E). In a separate set of Sim1^Cre^ animals, we injected the same AAV-Cre-mCherry vector into the LEC in combination with an AAV-Cre-GFP vector into the MEC. This approach enabled visualisation of the additional fan-cell projection to the MEC^60^ (Figure 4F).

**Figure 4.**
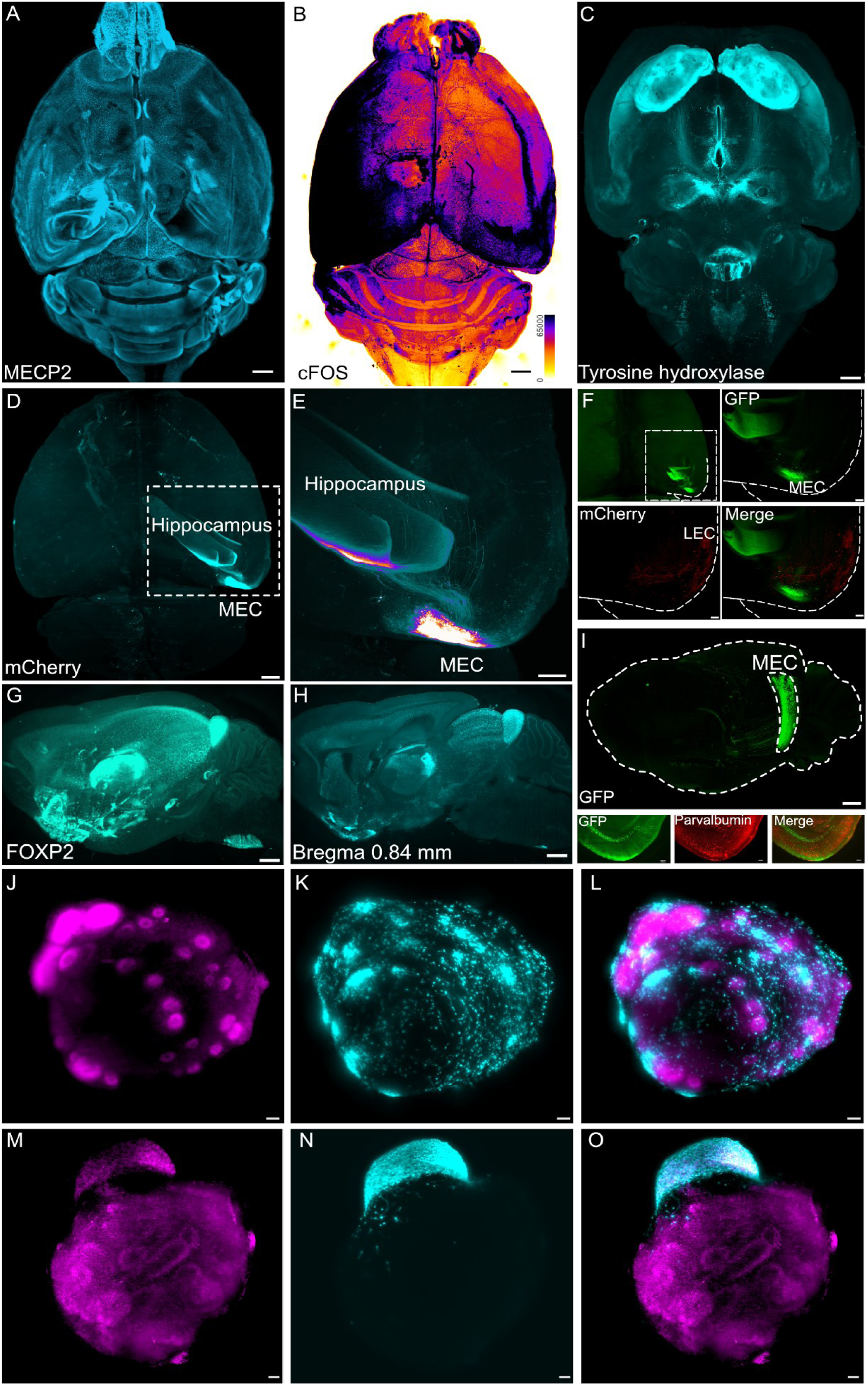
RatDISCO enables immunodetection of neuronal markers, viral tracing, and transgenic expression in mice and human brain organoids. Light-sheet microscopy images of whole-mouse brains immunolabelled for endogenous neuronal markers: MECP2 (A), cFOS (B, visualized with Inferno LUT), tyrosine hydroxylase (C), and FoxP2 (G: whole brain MIP; H-Optical slices at 0.84mm lateral to Bregma). D-F. Immunolabelling against virally-labelled medial (MEC) and lateral entorhinal cortex (LEC) neurons and axons in a Sim1^Cre^ mouse. (D) Whole-brain view of MEC projections. (E) Higher-magnification view of MEC-hippocampal projections. (F) Whole-brain view and a close-up of virally labelled MEC (GFP, green), LEC (mCherry, red) projections and merged image. (I) Immunolabelling of GFP in the transgenic P038 mouse line, highlighting MEC expression. Insets show GFP (green) co-labelling with parvalbumin (red) and the merged signal in a horizontal view. (J–O) Human iPSC-derived brain organoids (J, M; TO-PRO, magenta) co-cultured with GFP-positive single tumour cells (K, cyan) or tumour spheroids (N, cyan), with merged images shown in (L, O). Scale bars A-D, G-H: 500 µm. E: 250 µm, F, J-O and I-insets: 100 µm.

We also applied RatDISCO to enable detection of immunolabelled neurons and axonal projections in the enhancer trap PiggyBac transgenic mouse line P038^61^ (Figure 4I). This line expresses mCitrine in deep layers of the MEC, and we preserved these mCitrine-expressing neurons with an anti-GFP antibody. Whole-brain immunolabelling using RatDISCO revealed previously unreported mCitrine-expressing neurons in the subiculum and the retrosplenial cortex (RSC). Axonal projections from these neurons could be traced rostrally, surrounding the hippocampal formation and extending to the thalamus. To confirm the identity of these cells, we performed double immunolabeling against GFP and parvalbumin (Figure 4 I-insets horizontal view) and verified that the GFP-positive neurons are located in layer 5 of the MEC.

While RatDISCO is effective for large tissue samples, we also tested its compatibility with smaller models, such as organoids. Specifically, we applied RatDISCO to induced pluripotent cells (iPSC)-derived brain organoids co-cultured with GFP-labelled human glioblastoma (GBM) cells, the most common and aggressive primary brain tumour. Organoids (magenta; Figure 4 J-O) were co-cultured either with GFP-positive single tumour cells (cyan; Figure 4K) or with tumour spheroids (cyan; Figure 4N). We observed that single tumour cells distributed broadly across the organoid surface (Figure 4L, merged image), while tumour spheroids remained largely localised at the surface, showing minimal or no infiltration into the organoid interior (Figure 4O, merged image). Visualising these interactions between tumour cells and microenvironment can aid to understand metastasis progression.

These findings demonstrate that RatDISCO is broadly applicable, enabling whole-brain labelling and imaging in rodent models, including transgenic lines and viral tracing experiments, as well as in human-derived organoids.

### RatDISCO enables whole-amygdala mapping of stimulus-evoked neuronal activation in WT and Fmr1^-/y^ rats

To determine whether RatDISCO can identify behaviour-related circuit alterations, we compared cFOS expression, an immediate-early gene used as a proxy for neuronal activation, in the amygdala of WT and *Fmr1^-/y^* rats exposed to different elements of a classical fear conditioning paradigm. Naïve rats were compared to rats that experienced either a light stimulus (LS) alone or LS paired with a mild foot shock (FS; Figure 5 A-F). Using our custom 3D amygdala atlas and analysis pipeline, we quantified cFOS-positive cell density across the entire amygdala (Figure 5 G).

**Figure 5.**
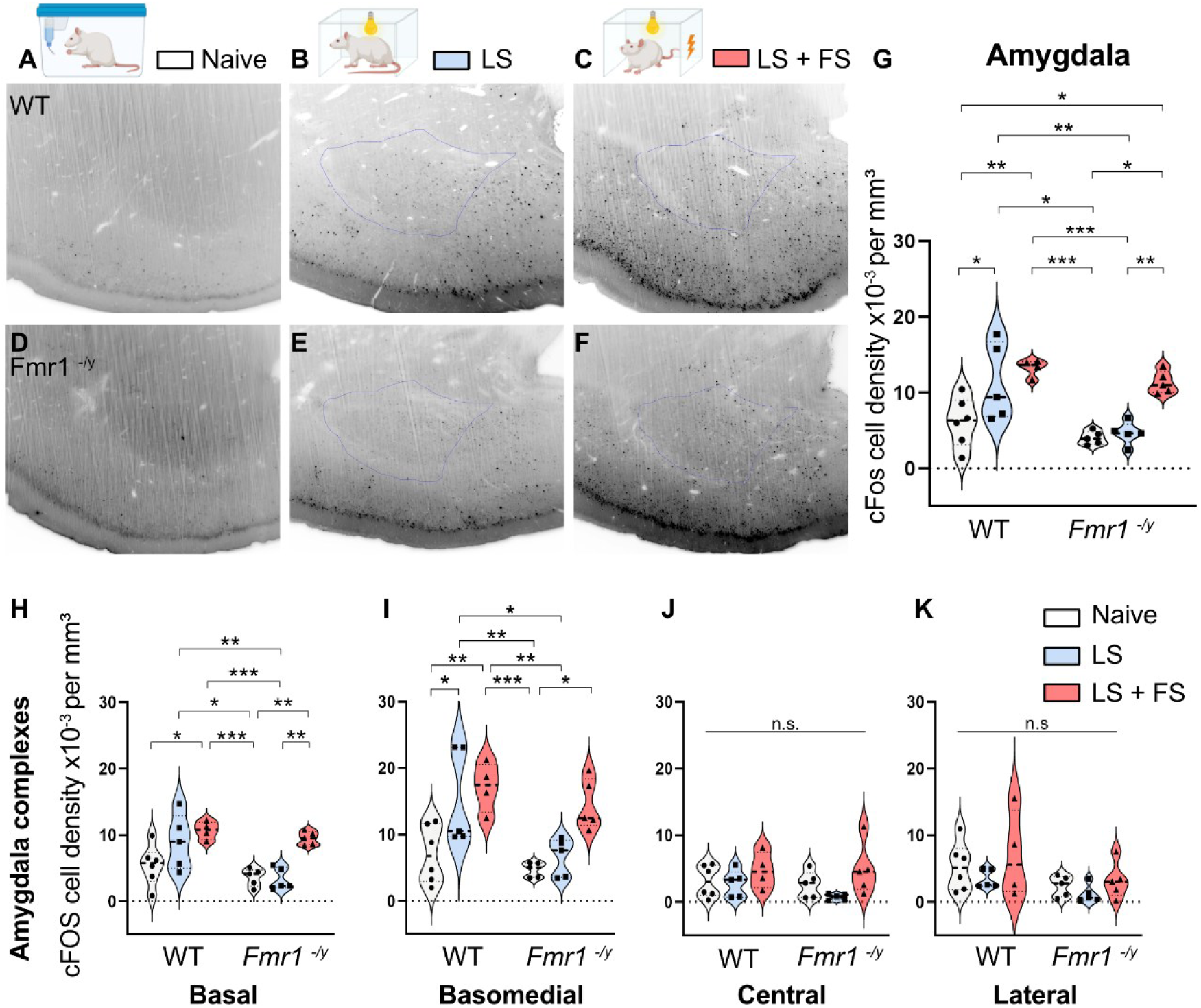
RatDISCO enables the quantification of experience-dependent cFOS expressing neurons in the amygdala of WT and *Fmr1^-/y^* rats in response to stimuli. Representative optical slice images of cFOS immunolabelling (black) in the amygdala of naïve rats (A, D, white), LS-exposed rats (B, E, blue), and LS+FS-exposed rats (C, F, red). G-K. Violin plots showing the total quantification of cFOS-positive cell density, expressed as x10^-3^ per mm³. In the entire amygdala (G), Circles represent naive, squares represent LS-and triangles represent LS+FS-exposed individual animals. Dashed lines inside the violin plots show median and quartile values. n.s > 0.05; *P < 0.05; **P < 0.01; *** P < 0.001. Tukey’s multiple comparisons and individual statistical values are described in supplementary Tables 3-6.

Significant differences in cFOS-positive cell density were observed across genotypes and experimental conditions (naïve, LS and LS+FS). WT animals showed higher cFOS-positive cell density than *Fmr1^-/y^* rats overall (2-way ANOVA: F_genotype_ (1, 24) = 12, P = 0.0017). In WT rats, exposure to LS and LS+FS significantly increased cFos-positive cell density compared to the naïve condition, suggesting sensitivity to the light stimulus. In *Fmr1^-/y^* rats, no significant difference was observed between naïve and LS conditions, suggesting a blunted response to the light alone. However, cFos cell density was significantly higher in Fmr1-/y following LS+FS rats, and in this condition, cFos-positive cell densities were comparable to those of WT rats (Figure 5 G, Tukey multiple comparisons and Supplementary Table 3).

These results show that WT rats expressed higher cFos in response to LS stimuli alone compared to *Fmr1^-/y^*rats, whereas both genotypes showed similar increases in cFos expression when compared naive animals to LS-FS experienced condition.

To determine whether these activation differences were uniform across the amygdala, we applied our 3D atlas comprising the basal, basomedial, lateral, and central complexes, and their subdivisions^25^. We then carried out the quantification of cFOS-positive cell densities within each subregion across genotypes and conditions (Figure 5 H-K; Supplementary Figure 3 and supplementary tables 3-16).

The cFos expression pattern observed in the whole amygdala was recapitulated in the basal and basomedial complexes (Figure 5H/I, respectively), but not in the lateral or central regions (Figure 5J/K, respectively), indicating that these two subregions predominantly drive the overall amygdala response. Together, these results demonstrate that RatDISCO can sensitively detect region-specific alterations in experience-dependent neuronal activation in the rat amygdala and highlight its potential as a powerful tool for mapping circuit-level differences in rat models of disorders.

## Discussion

In developing RatDISCO, we integrated traditional antigen retrieval with modern solvent-based clearing to create a robust, accessible protocol optimised for whole-brain immunolabelling in rats. Unlike other complex approaches, RatDISCO relies on commercially available antibodies and avoids the need for custom nanobodies or signal amplification, making it cost-effective and broadly applicable. RatDISCO is specifically tailored to the anatomical scale of rat brains, which are underrepresented in tissue-clearing development, while remaining compatible with mouse tissue and cerebral organoids. By combining high-temperature, alkaline antigen retrieval with staged delipidation and refractive index matching, RatDISCO enhances antibody penetration and tissue transparency without compromising structural integrity. It enables multiplex labelling and quantification of diverse neuronal populations, microglia, and vasculature in adult rat brains, and is compatible with viral tracers to study anatomical connectivity using light-sheet microscopy. RatDISCO enables brain-wide mapping of activity-dependent markers such as cFOS^19–21,63–67^, providing a powerful tool to study functional circuit alterations in animal and organoid models. This approach offers valuable insight into the neural mechanisms underlying neurodevelopmental disorders.

To evaluate RatDISCO against established tissue clearing protocols, we compared its performance with other common approaches. While mouse-tailored iDISCO+ resulted in optically transparent rat tissue, other approaches like TDE or ECi did not (Figure 1). We chose iDISCO+ as a comparison method due to its reported immunolabelling advantages in immunolabelling over other DISCO-based methods^23^. For this comparison, we focused on the transcription factor FoxP2, previously shown to be compatible with iDISCO+ and known for its distinct, spherical nuclear staining, an ideal feature for automated detection using machine learning algorithms and commercial image analysis software^71,72^.

Recent advances like iDISCO+ paired with ClearMap^16^ have enables quantification of nuclear markers by registering images to annotated atlases, primarily in mice. To extend this to rats, we developed a custom 3D amygdala atlas using Arivis Vision 4D, based on Paxinos and Watson’s Rat Brain Atlas, enabling accurate quantification of immunolabelled nuclei in light-sheet datasets. Compared to iDISCO+, RatDISCO showed improved antibody penetration and labelling compared under identical conditions. While our approach fills a gap in available tools, emerging registration tools such as QUINT and the Waxholm Space 3D rat brain atlas^73–77^ provide invaluable resources for future brain-wide analyses in cleared rat tissue.

In addition to enhanced labelling, RatDISCO provides superior signal in the far-red and near-infrared spectrum due to its higher light transmittance at longer wavelengths than iDISCO+. In contrast, iDISCO+ performed better at shorter wavelengths (e.g., 488 nm). These spectral properties suggest that for multiplex studies, RatDISCO immunolabelling is particularly well-suited for use with far-red fluorophores.

### Technical considerations with other clearing methods

Over 100 hydrogel-embedding, hydrophobic and hydrophilic clearing methods are available today^1,23–25^. All of these methods have their advantages and disadvantages. Hydrophilic methods (e.g., using urea or sorbitol) offer biocompatibility^78,79^, while hydrogel-based methods preserve endogenous fluorescence but limit structural integrity and long-term tissue re-imaging^80–84^. Solvent-based methods like RatDISCO rely on dehydration, immunolabelling, delipidation, and refractive index matching. These processes eliminate endogenous fluorescence, making robust antibody labelling essential. RatDISCO’s improved immunolabelling capabilities and versatility make it ideal for identifying distinct neuronal subtypes, glial cells and vasculature across various experimental systems and species.

RatDISCO can take 3–4 weeks in total due to the longer incubation times required for large rat brains in comparison to mice. This is longer than other methods like FAST 3D Clear (3 days for mice^85^) or traditional histology protocols. However, RatDISCO antibody incubation process does not require constant supervision and still offers a significant reduction in time and labour required for sectioning, labelling and imaging a whole brain into thin slices.

### Limitations of RatDISCO

RatDISCO leads to a 25% tissue shrinkage 25%, less than 3DISCO (∼50%) but more than iDISCO+ (∼11%). iDISCO+ uses milder solvents like DCM/methanol to limit shrinkage, avoiding the hazards of chloroform used in the original Folch lipid extraction method for biological samples^33,86,87^. Unfortunately, methanol is a milder solvent for lipid extraction compared to THF and requires extended incubation times to work effectively on mouse brains^88^. Still, these solvents compromise lipid extraction efficiency in large rat tissue and results in brittle tissue. Other methods, like uDISCO^89^, achieve further clearing using tert-butanol and BABB but lead to greater shrinkage (∼35%) and pose higher toxicity risks, complicating handling and imaging.

RatDISCO uses SDS as part of the antigen retrieval pre-steps. SDS is commonly used in water-based clearing methods to form micelles that remove the lipids in the cell membrane^11,81,82,90^. However, prolonged SDS incubation without hydrogel scaffolding compromises tissue integrity, leading to expansion or collapse. For this reason, we recommend limiting antigen retrieval for short periods, ideally 2 days at 37°C or 1 hr at 80°C.

A practical challenge has been the inconsistent quality and availability of THF, which is critical for tissue dehydration and transparency. Batch variability, sometimes within the same brand, can disrupt clearing. We tested several THF variants (Supplementary Table 12) and observed significant inconsistencies in the quality of THF batches. Nevertheless, we have identified two reliable THF sources that work with our protocol. It is essential to use anhydrous THF with added butylated hydroxytoluene (BHT) as an antioxidant and stabiliser to achieve high transparency. Finally, greener alternatives like 2-methyl-THF may offer safer, more sustainable options for future adaptations that could be further implemented.

Overall, RatDISCO is a versatile and robust tissue-clearing and immunolabelling protocol that enables comprehensive labelling of neuronal subtypes, glia, and vasculature across brain regions. Its broad applicability across species makes it a powerful tool for studying complex neuroanatomy and uncovering circuit-level alterations in models of neurodevelopmental disorders.

## Acknowledgements

This work was funded by the Simons Initiative for the Developing Brain. DS and PS are funded by Cancer Research UK with a Programme Foundation Award to DS (DRCPFA-May21\100011). We sincerely thank Professor Matthew Nolan for his guidance and invaluable feedback throughout this project. We are also grateful to Dr. Crispin Jordan for his insightful advice on statistical approaches. We acknowledge ST for conducting all behavioural experiments and BV for performing the viral experiments in Sim1^Cre^ mice. AGS performed intrahippocampal kainate injections, while PS conducted the organoid experiments under DS guidance. NR, PK, and ID provided invaluable guidance and feedback throughout the project. BG processed tissue for Figure 2. KJC and CMG were responsible for all tissue processing, imaging and quantification; CMG carried out the analysis and wrote the manuscript, to which all authors contributed.

## Materials and methods

### Animals

All animal experiments were approved by the University of Edinburgh veterinary services and performed per the guidelines established by European Community Council Directive 2010/63/EU (22 September 2010) and the Animal Care (Scientific Procedures) Act 1986. All animal experiments were approved and conducted following the University of Edinburgh Animal Welfare Committee, in accordance with the guidelines established by European Community Council Directive 2010/63/EU (22 September 2010) and by the Animal Care (Scientific Procedures) Act 1986, and by the Animal Care (Scientific Procedures) Act 1986. All animals were kept on a 12-hour light/dark cycle and had ad libitum access to water and food.

Animals were anaesthetised with isoflurane followed by a lethal dose of sodium pentobarbital (Euthatal), then transcardially perfused with phosphate-buffered saline (PBS; Invitrogen) followed by freshly made 4% paraformaldehyde (PFA; Sigma Aldrich) in 0.1 M phosphate buffer (PB; Sigma Aldrich). Brains were removed and post-fixed for 3 hrs in 4% PFA, then rinsed in PBS. The tissue was then stored in PBS with sodium azide 0.05% (Sigma-Aldrich 71289) at 4°C until it was used.

### Rats

Male LE-*Fmr1^em1/PWC^*, hereafter referred to as *Fmr1*^-/y^, and WT littermates, bred in-house and kept in a 12-hour/12-hour light/dark cycle with ad libitum access to water and food, were used. Colony founders were produced by SAGE (Sigma Advanced Genetic Engineering) Labs using ZFN-mediated disruption of *Fmr1^-/y^* with a targeted construct containing the coding sequence for eGFP; resulting founders did not express FMRP or eGFP. Pups were weaned off their dams at P22 and housed in mixed-genotype cages with littermates (three to four animals per cage).

### Mice

C57bl/6J mice aged 5-12 weeks were obtained from Charles River UK or bred in-house. P038 mice aged 8-16 weeks^40^. Sim1^Cre^ mice (generated by GenSat and obtained from MMRRC: Tg(Sim1cre) KH21Gsat/Mmucd) were bred to be heterozygous for the Cre transgene by crossing male mice carrying the transgene with female C57BL6/J mice (Charles River, UK). Experiments labelling axonal projections from MEC/LEC to the hippocampus used Sim1^Cre^ mice aged 7-14 weeks.

### Acute intrahippocampal kainate temporal lobe epilepsy mice model

To generate acute seizures and cFOS activation, mice were anaesthetised with isoflurane and were injected with 1 ml of 5% dextrose saline, to prevent dehydration during status epilepticus. Kainate (100 nl, 20 mM in saline, Tocris) was infused into the left dorsal hippocampus targeting the molecular layer of the dentate gyrus (stereotaxic injections coordinates: 1.85 mm caudal, 1.25 mm lateral from bregma and 1.40 mm ventral from brain surface). Mice were allowed to recover and were perfused 1.5 hours after the occurrence of the first generalised seizure.

### Viral labelling of axonal projections in mice

For injections into the MEC in Sim1^Cre^ mice, a craniotomy was made between the lambdoid suture and sinus at 3.50 mm lateral to midline and three injections were made of adeno-associated virus with a pipette angled backwards at 6°, 8° and 10°. For injections into the LEC, a craniotomy adjacent to the intersection of the lambdoid suture and the ridge of the parietal bone was made approximately 3.8 mm posterior to bregma and 4.0 mm lateral to bregma with a pipette angled laterally at 12°, 10°, and 8°. We used a rAAV2-eF1a-DIO-mCherry, titer: 2.6×10^-12^ GC/ml (Addgene #: 114472) or a AAV1/2-FLEX-GFP (Addgene #: 28304) titer: 2.44×10^13^ GC/ml, generated in-house. We used 150-200 nL per injection. For all injections, a glass pipette was slowly lowered into the brain until a slight bend indicated an approach to dura. The pipette was then retracted 200 uM and the virus was injected at that location. The pipette was retracted after a stationary interval of four minutes, the incision was closed with sutures, and mice were administered an oral analgesic prepared in flavoured jelly immediately after surgery. Mice were perfused 3-4 weeks after surgery.

### iPSCs cell culture

Human Nas2 iPSCs^91^ were maintained in StemMACS iPSC brew XF medium (Miltenyi Biotec) using a feeder-free culture protocol in six-well plates (Corning) coated with laminin 521 (Biolamina). iPSCs were cultured at 37°C and 5% CO_2_ with daily feeding of 2mL media per well. Cultures were grown to no higher than 80% confluency. Passaging of iPSC colonies was carried out using EDTA (Life Technologies) to the desired split ratio. Human Nas2 iPSC line was a kind gift from Prof. Tilo Kunath.

### Human glioblastoma (GBM) cell culture

GMB cell lines E17 were grown on T25 or T75 flasks (Corning) coated with laminin (Cultrex) in serum free media at 37°C, 5% CO_2_. Media composition as follow: DMEM/F12 (Merck), Glucose (Merck), MEM-NEAA (Gibco), Pen/Strep (Gibco), BSA solution (Gibco), bMercaptoEtOH (Gibco), B27 and N2 supplement (Gibco), Laminin (Cultrex), mouse EGF (peprotech) and human FGF (peprotech). When reaching 70-75% confluency, cells were lifted using Accutase (Innovative Cell Technologies), washed with DMEM/F12 (Merck), Pen/Strep (Gibco), BSA solution (Gibco), then resuspended in fresh media and plated on Laminin coated flasks.

### iPSC organoids

Organoids were generated as previously described^92^ with minor modifications. In brief, when iPSC cultures reached ∼70% confluency, the medium was aspirated and wells rinsed twice with DMEM/F12.1mL of Accutase (Innovative Cell Technologies) was added per 6-well and incubated for ∼5 minutes at 37°C, 5% CO_2_ until cells detached from the dish. Using a 1000μL pipette, gentle trituration was performed to achieve a single cell suspension, which was transferred to a 50 mL conical tube. Cells were washed with DMEM/F12 three times. Cells were counted using a haemocytometer and resuspended at approximately 100,000 cells/mL in E8 medium (Life Technologies) plus 10 μM ROCK inhibitor Y-27632 (Tocris). 10,000 cells/well were seeded into 96-well V-bottom low-attachment plates (Corning). At 24 hours (day 0) media was replaced with E6 medium supplemented with 2.5 μM dorsomorphin, 10μM SB431542, and 2.5 μM XAV-939 (Cayman chemicals); 100 μl of this media was added to each well and a further 100 μl of media added at day 1. Wells were fed daily with ∼50% medium exchange from day two until day five. On the sixth day, the medium was changed to neural medium (NM), consisting of Neurobasal-A plus B-27 supplement without vitamin A, GlutaMax and Anti-A and supplemented with 20 ng/mL EGF plus 20 ng/mL FGF2 (peprotech). NM plus EGF/FGF2 (Medium B) was changed daily for 19 days with ∼75% media exchanges. Spheroids were transferred to 6-well low-attachment plates (Corning) on day 25. To promote differentiation of neural progenitors into neurons, FGF2/EGF were replaced with 20 ng/mL BDNF and 20 ng/mL NT3 (Peprotech) from days 25-43 and media changed every other day with ∼75% media exchanges. From day 43 onward, medium changes were done three days per week using neural medium without added growth factors. Throughout the culture period, organoids that fused together were separated by cutting with a disposable scalpel (Swann morton).

### Organoid-GBM spheroid assembloids/co-culture

E17pbGFP cells were detached using accutase, counted, and 100,000 GBM cells seeded in 200 µl of complete media per well of 96-well V-bottom low-attachment plate (Corning) with daily full media changes. At day seven, one ∼80-day old organoid was added to each well using a Pasteur pipette and plates were spun at ∼100 g before a complete media change to neural medium without added growth factors. Brain organoid-GBM spheroid assembloids were cultured for a further seven days with daily complete media changes. Finally, tissue was immersed in 4%PFA at 4℃ for 3 hrs before RatDISCO processing.

### Light-sheet microscopy imaging

Cleared samples were imaged in the sagittal, horizontal or coronal plane using a Gaussian beam light-sheet microscope (Ultramicroscope II or Ultramicroscope Blaze LaVision-Miltenyi Biotec). The Ultramicroscope II was equipped with an Andor Neo sCMOS camera and Olympus MVPLAPO 2x objective with 6 mm working distance (WD) dipping cap adjusted to 2.152x. Filter sets: Ex 488 nm - Em 525/50 nm; Ex 561 nm - Em 595/40 nm; Ex 640 nm - Em 680/30 nm. The Blaze was equipped with a sCMOS camera 4.2 Megapixel, Light Speed Acquisition model and LaVision-Miltenyi BioTec objectives: 1.1x/0.1 NA MI PLAN objective (WD 17 mm), 4X/0.35 NA MI PLAN objective and 12x/0.53 NA MI PLAN objective (WD 10 mm) with dipping caps for organic solvents. The BLAZE was equipped with a laser beam combiner with following laser modules including the Ø43mm emission filters: Ex 405 nm - EMF460/40m; Ex 488 nm - EMF525/50m; Ex 561 nm - EMF620/60m; Ex 640 nm - EMF680/30m and Ex 785 nm - EMF805LP.

The tissue was imaged with a step size between 4 - 10 µm using double side illumination and contrast blending algorithm from the IMspector acquisition software (Miltenyi Biotec). Images were adjusted for brightness and contrast in FIJI and analysed using QUINT or Arivis Vision 4D software.

### Image analysis

cFos and FoxP2 immunolabelled neurons were quantified in Arivis Vision 4D (now Zeiss Arivis Pro) software. First, we created a template atlas and an analysis pipeline, which were applied to all image data sets.

We aligned the low-magnification images of the entire brain in the sagittal plane to the higher-magnification images of the amygdala. We then manually created a 3D template atlas based on anatomical landmarks in the tissue corresponding to levels 3.4 mm - 4.6 mm lateral to Bregma (The Rat Brain Atlas in Stereotaxic Coordinates Third Edition, Franklin and Paxinos; Figures 175-180).

### 3D amygdala atlas

We manually registered whole-brain, low-magnification light-sheet images to the Paxinos and Watson atlas in FIJI and Arivis Vision 4D. We used major anatomical landmarks to register images corresponding to Bregma 3.4 mm - 4.6 mm to create independent 2D templates per anatomical plane (Paxinos and Watson figures 175-180). We then aligned these to high-magnification light-sheet images of the amygdala to create a volumetric 3D atlas of the rat amygdala using Arivis. We then used FIJI to adjust the size in the x and y dimensions of the Franklin and Paxinos atlas images to match the corresponding level for the light-sheet microscopy images. Subsequently, we imported the full data sets into Arivis. In Arivis, we used the “draw objects” tool to create new “objects” or ROIs in the shape of each amygdala complex and subnuclei and followed them through its z-depth, thus creating a 3D atlas for our structure of interest. We saved these “3D objects” as a template atlas and imported them to all analysed image data sets. We aligned this 3D atlas in x,y and z to match the anatomical landmarks on each animal before running the analysis pipeline.

The amygdala atlas subdivisions^25^ included the basal complex, which includes the basolateral anterior (BLA), posterior (BLP), and ventral (BLV) nuclei. The basomedial complex that includes the posterior (BMP) and anterior (BMA) nuclei. The lateral complex that includes the dorsal-lateral (LaDL) and ventromedial (LaVM) nuclei. And the central complex that includes the lateral (CeL), medial (CeM), and central (CeC) nuclei and the extension of the amygdala (EA).

### Analysis pipeline

Using our 3D amygdala template atlas, we established an analysis pipeline for automatic cell detection using Arivis Vision 4D. Click on the “analysis” menu, select “analysis panel”, and create a new “analysis pipeline”. On the Input ROI menu, select the bounds (all), planes (number of images in z), time points (1), image set (file name/location), and channels (according to fluorophore used). Add “blob finder” as a new operation in the analysis pipeline.

In blob finder; Input ROI menu: Select the “channel” of interest. Measure and set the diameter of your objects of interest using the “ruler” tool. The average size of cFOS cell nuclei in our images was 12 µm in diameter. Adjust the “threshold” to find the optimal results for segmenting your objects of interest. We refined it to 6-8% in 6 animals and used that value for all. Finally, adjust the “splitting sensitivity” to find the optimal segmentation when two or more cells are near each other, resulting in 70%.

Add “import document objects” as a new operation in the analysis pipeline. In import document objects: Add objects assigned with a “tag”. See “create an atlas”. Fill up the provided text box for all available tags. Selected tags will be included as output for all subsequent operations in the analysis pipeline: Check the option “include compartments” to import “children” (“children” are blobs or spherical cell nuclei detected within an object). Check the option “filtered by tag” to select additional outputs. This option is useful when you want to load a subset of a particular object group, such as nuclei within “objects”. In this case, you would use “ECIC” or “CIC” in tags, check to Include compartments and use “nuclei” filtered by tag. Export and save the newly created “analysis pipeline”.

Once the analysis pipeline is created, it can be imported and used for all experimental animals.

### Validation and manual cell quantification

Automatic quantification was done using the same threshold, split sensitivity, and cell soma size parameters for all animals. These parameters were used to detect all cFOS-positive nuclei in each major basolateral complex of the amygdala from Bregma 3.4 mm—4.6 mm. They were manually validated across 12 animals before being applied to all analysed images (Supplementary Figure 2).

Manual quantification was performed using the CellCount plugin in FIJI. Using our high magnification images (3.2X), we identified the starting plane of the amygdala (atlas figure 175) within each of our animals. We imported each ROI set to the corresponding images within the data stack using the 2D template atlas that we delineated for each anatomical plane (Figures 175 - 180). Once the ROIs were imported, we duplicated the image and flattened it with all ROIs visible. We then manually counted all cFOS-positive cells present in each ROI. This was done for 1 in every 10 images at each anatomical level across 12 animals.

Total cell counts were calculated per region and used to refine our automatic cell quantification parameters. The total number of cells manually counted was n=44,682. The total number of cells manually counted per basolateral complex was as follows: basal complex n=15,255; basomedial complex n=16,602; central complex n=6,082; and lateral complex n=1,926.

For each subsequent set of images, follow the next steps: Open a new set of images, import the corresponding “object”/template atlas of the IC and adjust it in x,y, and z using the “move objects” tool. Import the saved “analysis pipeline” and run it. Export your measurements to Excel or via CSV for further statistical analysis. Our amygdala template atlas and Arivis analysis pipeline are available upon request.

### Behaviour

Male WT and *Fmr1^-/y^* littermate rats were acclimated to a holding room and handled for 5 min/day for two days before habituation to a neutral context (a modified Coulbourne Instruments Habitest box dimensions 30 cm × 25 cm × 32 cm containing a curved black and white striped wall insert, smooth grey floor and no electrified grid) for two 5 min sessions on non-consecutive days (2 or 3 days apart). Conditioning followed on the day after the second test context habituation and was performed in a standard, unmodified Habitest box, including an electrified grid floor and cleaned with disinfectant wipes (Distel). Conditioning occurred over 21 minutes and consisted of 3 minutes to explore the conditioning context, followed by 6 pairings of a flashing light stimulus (LS) co-terminating with a mild foot shock (FS). The LS was a 10 sec flashing light (5Hz maximum intensity flashes, 50/50 duty cycle); the FS was a 1 sec, 0.8 mA scrambled shock delivered through the bars of the floor; LS presentations started at 180, 360, 490, 770, 980, and 1280 sec into the training period. Controls included rats that received 6 presentations of the flashing light (LS), but did not experience the FS during training and those that did not undergo any form of training (naive).

All behavioural experiments were carried out during the light phase, between 08:00 and 10:00 hrs, to maintain consistency in the circadian rhythms that could affect cFOS expression.

### Shrinkage

For comparison, hemispheres from the same animal were cleared using either RatDISCO (right side) and iDISCO+ (left side); or TDE (right side) and ECi (left side). We measure the distance in millimetres in the anterioposterior, mediolateral and dorsoventral axes of 4 rat brains, excluding the olfactory bulbs and the spinal cord, before and after clearing using FIJI.

### Light transmittance

The light transmittance of tissue cleared with RatDISCO and iDISCO+ was measured with a spectrophotometer Ultrospec 2000 (Pharmacia Biotech). For comparison, hemispheres from the same animal were cleared using either RatDISCO (right side) or iDISCO+ (left side). The hemispheres were placed in a cuvette (Fisher Scientific, 15520814) filled with refractive index matching solution ECi. The transmittance of the light was measured from 400-850 nm in 20 nm increments for all samples. Beams were orientated perpendicular to the sample and passed through the central part of the tissue. All samples were placed following the same anatomical orientation towards the beam. The reference or blank values were obtained using a cuvette filled with ECi per wavelength before adding the samples. TDE-and ECi-cleared tissues were not measured because they were completely opaque.

### Detailed RatDISCO protocol

Antigen retrieval pre-treatment: Incubate the tissue in an antigen retrieval solution at 80°C in a water bath or similar for 1 hr (4% SDS and boric acid 2.8 mM in dH2O; adjust the pH to 8.5 using NaOH 10M). We do not recommend using a microwave because of hotspots that can damage the tissue. Let the samples cool down at RT for 30 min. Finally, wash the tissue several times in PBS to remove the SDS.

Dehydration MeOH/PBS dehydration/rehydration and permeabilization: Incubate the tissue in a series of methanol-PBS steps (table below). Then, wash the tissue 3 times for 30 min in PBS at RT. All MeOH dilutions are made with PBS.

| *MeOH/PBS* | *Incubation time* | *Incubation time* | *Incubation time* |
| --- | --- | --- | --- |
|  | **Rat** | **Mouse** | **Organoid** |
| 50% | 1.5 hrs | 1 hr | 0.5 hrs |
| 80% | 1.5 hrs | 1 hr | 0.5 hrs |
| 100% | Overnight | Overnight | 1 hr or overnight |
| 100% | 1.5 hrs | 1 hr | 0.5 hrs |
| 100% | 1.5 hrs | 1 hr | 0.5 hrs |
| 80% | 1.5 hrs | 1 hr | 0.5 hrs |
| 50% | 1.5 hrs | 1 hr | 0.5 hrs |

Blocking: Incubate the tissue in PBSGT blocking solution (0.2% gelatine, 0.01% sodium azide, 0.5% Triton X-100 in dH2O) under agitation at RT for 12-24 hrs.

Immunolabelling.

Primary antibody incubation: Incubate the tissue in primary antibodies at 37°C in PBSGT + 0.1% Saponin for 7-10 days (rats), 5-7 days (mice), or 1-2 days (organoids) under gentle agitation. Use enough antibody solution to cover the tissue completely and refresh the solution halfway through incubation. Use a small vial, bijoux or Eppendorf tube suitable in size for your tissue and cover the tissue completely with the antibody solution (i.e. 3-5 ml in 7 ml bijoux containing half brain hemisphere of a 2-month-old rat). Note: If your primary antibody is conjugated to a fluorophore, do not carry out secondary antibody incubation and proceed directly to the delipidation steps.

Secondary antibody incubation: Wash the tissue in PBS at RT every hour for 1 day and leave it in PBS overnight. Incubate the tissue in the secondary antibodies at 37°C in filtered PBSGT + 0.1% Saponin (see notes) for 5-7 days (rats and mice) or 1-2 days (organoids) under agitation.

Use enough antibody solution to cover the tissue completely and refresh the solution halfway through incubation.

Delipidation and optical clearing: Wash the tissue in PBS at RT every hour for 1 day and leave it in PBS overnight. Then, incubate your tissue in increasing concentrations of tetrahydrofuran (THF), followed by dichloromethane (DCM) and dibenzyl ether (DBE) as follows:

| *Solvent* | *Incubation time*<br>***Rat or Mouse*** | *Incubation time*<br>***Organoid*** |
| --- | --- | --- |
| *50% THF* | *1 hr* | *0.5 hrs* |
| *80% THF* | *1 hr* | *0.5 hrs* |
| *100% THF* | *1 hr* | *0.5 hrs* |
| *100% THF* | *Overnight* | *1 hr or overnight* |
| *100% THF* | *1 hr* | *0.5 hrs* |
| *100% DCM* | *30 min - 1 hr* | *15 min* |
| *100% DBE* | *1 hr* | *0.5 hrs* |

Imaging and storage: Transfer tissue to ethyl cinnamate (ECi) for imaging or long-term storage at RT. We recommend leaving the tissue to equilibrate in ECi overnight before imaging. All THF dilutions are made with water.

### Key points and tissue requirements

We recommend 4% paraformaldehyde (PFA) perfused-fixed tissue. Fixation and post-fixation times are critical for some antibodies. We recommend using freshly made 4% PFA for transcardial perfusions and between 3 and 24 hours of post-fixation at 4°C. Note that longer post-fixation times can be detrimental to immunolabelling. After post-fixation, tissue can be kept in PBS + Azide at 4°C for long periods of time (months) without noticeable effects on clearing or immunolabelling. However, we do not recommend using frozen tissue. Finally, we recommend using a large volume of all solutions before delipidation *i.e.,* 10 ml for a 15 ml falcon tube or 35 ml for a 50 ml falcon tube (steps 2-6). This is to facilitate the dehydration of the tissue and washes. Glass vials must be used for the delipidation and clearing (refractive-index matching; steps 6-7). We recommend filling the vials to the top to avoid oxidation and to facilitate delipidation.

### iDISCO+

Tissue was processed following the iDISCO+ protocol described by Renier *et al* 2016^16^.

### RNAscope adapted protocol for RatDISCO

Here, we summarise the steps for carrying out RNAscope in large rat brain hemispheres. We have outlined the approximate time needed to complete each step and highlight the possible stopping points within the protocol.

Tissue fixation and washing: After perfusion, tissue was post-fixed for 17–24 hrs in 4% PFA at 4°C. Tissue was then washed in 10 mL of 0.1 M PBS + 0.1% Tween-20 (PBS-Tween) for 1 hr at RT before sequential incubation in 20%, 50% and 75% MEOH in PBS-Tween (10 mL per sample for 1 hr at RT). Samples were processed immediately or stored in 100% MeOH at-20°C for up to two months (stopping point).

Hydrochloric acid (HCl) and MeOH pre-treatment (2.5 hrs): We incubated the tissue in 10 mL of 0.2 M HCl in 100% methanol at RT for 30 mins. We then incubated the tissue in 75%, 50% and 25% MeOH-PBS-Tween (10 mL per sample for 30 min each at RT) and then in 10 mL PBS-Tween + 1% BSA at RT for 10 mins.

Antigen retrieval (1 hr): We pre-warmed 25 mL of RNAscope 1X Target Retrieval solution in a water bath at 100°C before incubating the tissue for 15 min. We immediately transferred the tissue to a fresh falcon tube containing PBS-Tween + 1% BSA and incubated it at RT for 5 min, followed by a 5 min wash in 100% MeOH, and a 15 min wash in PBS-Tween + 1% BSA.

Protease treatment (40 mins): We decanted the PBS-Tween + 1% BSA and added 1 mL of RNAscope Protease Plus solution, and incubated the tissue for 30 min at 40°C (all incubations at 40°C were carried out in a water bath). We then decanted the protease solution and washed the sample for 5 min using enough PBS-Tween + 1% BSA or RNAscope probe diluent to cover the tissue (a possible stopping point is storing the tissue at 4°C for 1 day only).

Probe hybridisation (2.5 hrs): We diluted our probes (ACDBio 320881, 320871, 407821) according to the RNAscope protocol (C1: non-diluted; C2 and C3 diluted 1:50 in either C1 or RNAscope probe diluent reagent). Probes were warmed up at 40°C for 10 min before cooling at RT and then added to the tissue and hybridised for 2 hrs at 40°C. Tip: The temperature must be maintained at 40°C at all times. Tissue was washed with 1X RNAscope wash buffer (ACDBio 310091) 3 times for 10 min each at RT. (Optional stopping point: tissue can be stored in 5X SSC overnight for up to 7 days). Note: all subsequent “washing steps” were done in 1X RNAscope wash buffer 3 times for 5 min at RT.

Amplification and hybridisation (1 hr per amplification): For the sequential amplification of probes 1 and 2, tissue was incubated in RNAscope Multiplex FLv2 AMP1 or AMP2 at 40°C for 30 min before washing. For probe 3, tissue was incubated for 15 min RNAscope Multiplex FLv2 AMP3 before washing (ACDBio 323100).

Developing the signal (2 hrs per channel): We decanted the washing buffer, added RNAscope Multiplex FLv2 HRP-C1 solution to cover the sample, and incubated it at 40°C for 15 min before further washing. We then decanted the washing buffer and added 1 mL of Opal dye (690, 570 or 488 diluted 1:1000 in TSA buffer) before incubating at 40°C for 30 min, followed by washing. Finally, we decanted the buffer and added RNAscope Multiplex FLv2 HRP blocker to cover the sample. We incubated at 40°C for 15 minutes and repeated the washing steps (optional stopping point). We repeated the process to develop the other probes using RNAscope Multiplex FLv2 HRP-C2 and/or RNAscope Multiplex FLv2 HRP-C3 with the relevant Opal dye (optional stopping point).

Once RNAscope labelling was complete, tissue was immunolabelled and/or cleared using RatDISCO. Note: all RNAscope solutions were prepared fresh before the assay using sterile ddH_2_O in autoclaved containers. Aseptic conditions remained throughout.

### Statistics

Throughout the manuscript the statistical test is stated, along with p values, and test statistics. The researchers conducting the experiments and analysing the data were blinded to the genotype of the rats and the treatment group. Statistical analyses were performed using GraphPad (Prism 10.0). Statistical significance was assessed using either t-tests, one-way or 2-way ANOVA followed by Tukey multiple comparison analysis. Rats were used as an experimental unit throughout the manuscript. Results are presented as means ± SEM. Summary of statistics described in Supplementary Tables 1-11.

