## Supplementary material for "RatDISCO, a tissue clearing and immunolabelling protocol for large rat brains": Suplementary data

### Supplementary Figures and Tables

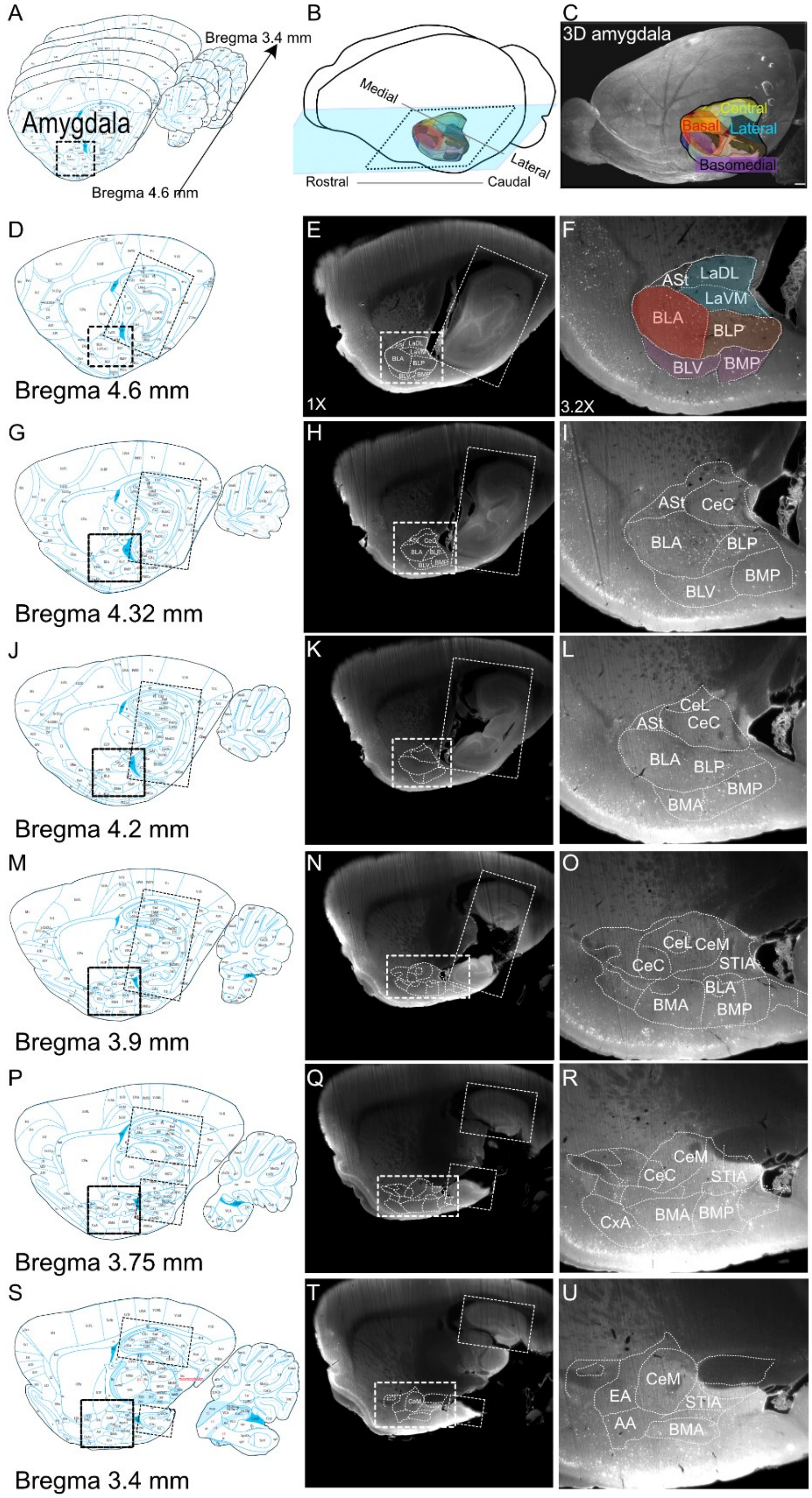

**Supplementary Figure 1. Registration of rat amygdala atlas to light-sheet microscopy images to create a 3D amygdala atlas.**

Schematic figures from the Paxinos and Watson atlas corresponding to the sagittal levels lateral to Bregma 4.6 mm, 4.32 mm, 4.2 mm, 3.9 mm, 3.75 mm, and 3.4 mm (A, D, G, J, M, P and S; Dotted box highlights the amygdala). B. Schematic figure of the 3D template amygdala atlas created in ARIVIS in relation to the brain. C. Light-sheet microscopy image of a rat brain with the registered 3D amygdala atlas.

E, H, K, N, Q, and T. Low magnification light-sheet microscopy representative images of a rat brain matching the corresponding anatomical level of the Paxinos and Watson atlas on the left and used to register the template amygdala atlas. White dotted insets highlight the anatomical landmarks used to align to the correct mediolateral levels. F, I, L, O, R, and U. Matched high-magnification images of the amygdala with the registered template atlas (dotted lines), including the amygdala's major complexes and subdivisions. The basolateral complex division (made up of the basal, basomedial and lateral cell groups; and the central nucleus (CeA). The basal complex (red) includes the basolateral anterior (BLA), posterior (BLP) and ventral (BLV). The basomedial complex (purple) includes the posterior (BMP) and anterior (BMA). The lateral complex (blue) includes the dorsal-lateral (LaDL) and ventro-medial (LaVM) nuclei. Additionally, the central complex (yellow) of the amygdala is subdivided into: lateral (CeL), medial (CeM) and central (CeC) nuclei.

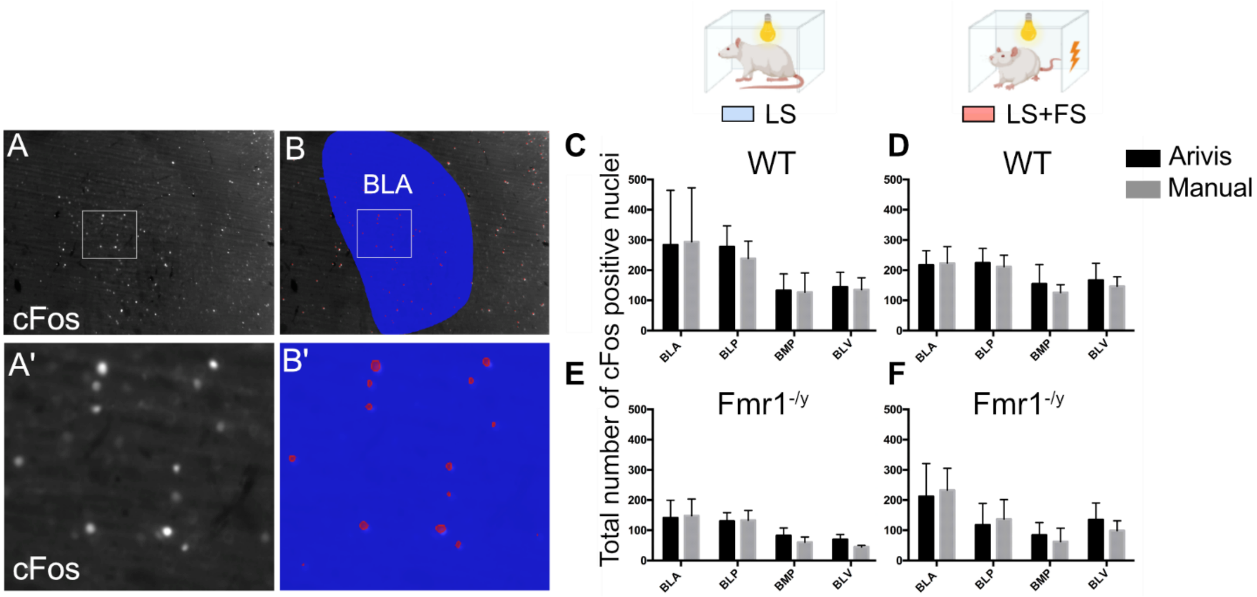

**Supplementary Figure 2. Manual vs automatic cFOS cell quantification validation.**

A. Light-sheet image of a rat brain immunolabelled against cFOS. B. Automatic cFOS-positive cell detection by Arivis in red on light-sheet images of the basolateral anterior amygdala (BLA in blue). Insets A' and B' show higher magnification of the images above. Total number of cFOS-positive neurons (nuclei) manually counted in grey versus automatically in black. C-F. Column graph bars showing the comparison between WT (C-D) and *Fmr1*$^{-/y}$ rats (E-F) exposed to light stimuli only: LS or LS and mild foot shock: LS+FS (mean ± SEM; n=3 animals per condition).

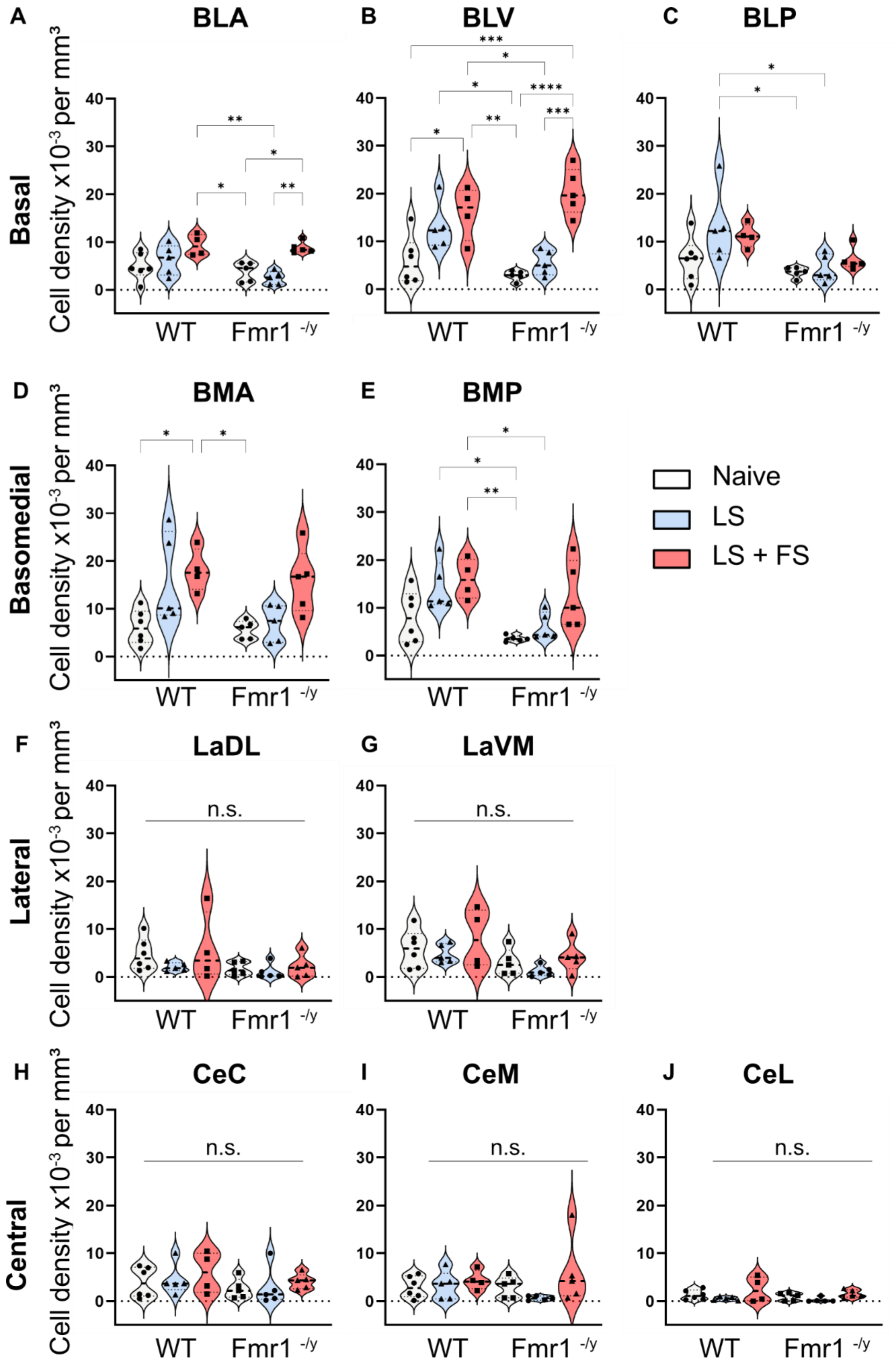

A
BLA
B
BLV
C
BLP
Basal
Cell density x10-3 per mm3
WT
Fmr1 -/y
D
BMA
E
BMP
Basomedial
Naive
LS
LS + FS
F
LaDL
G
LaVM
Lateral
n.s.
H
CeC
I
CeM
J
CeL
Central

**Supplementary Figure 3. Quantification of experience-dependent cFOS expressing neurons in the subnuclei of the amygdala.**

Violin plots showing the cFOS cell density $x10^{-3}$ per $mm^3$ in the amygdala complexes and its subnuclei of WT and *Fmr1*$^{-/y}$ rats exposed to naive, LS and LS+FS conditions. Basal complex containing the BLA (A), BLV (B) and BLP (C). Basomedial complex containing the BMA (D) and BMP (E). The Lateral complex containing the LaDL (F) and LaDM (G). The Central complex containing the CeC (H), CeM (I) and CeL (J). n.s > 0.05; *P < 0.05; **P < 0.01; *** P < 0.001. Tukey's multiple comparisons and individual statistical values are described in supplementary Tables 7-16.

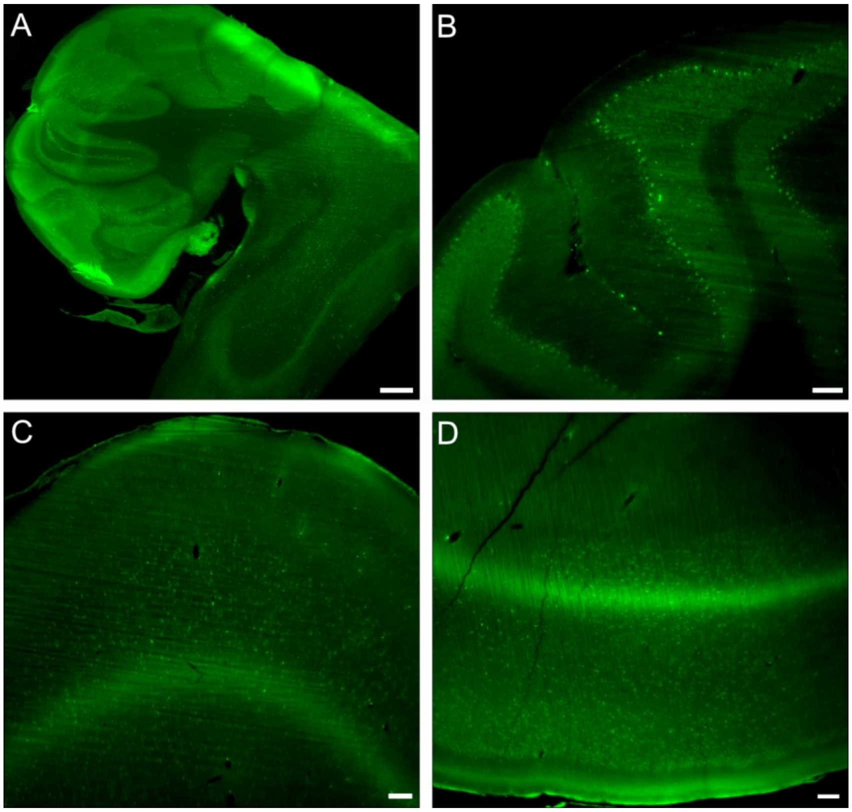

**Supplementary Figure 4. RatDISCO is compatible with RNAscope *in situ* hybridisation.** Light-sheet images of rat brains labelled with a POLR2A probe (green) using a modified RNAscope *in situ* hybridisation protocol then processed using RatDISCO tissue clearing. A. Maximum intensity projection of the cerebellum and brainstem regions. Single optical-slices of the cerebellum (B), the prefrontal cortex (C) and the amygdala area (D). Scale bar A: 500 µm, B-D: 100 µm.

| Wavelength (nm ) | RatDISCO (mean +/- SD) | iDISCO+ (mean +/- SD) | P value | t, df | P value summary | Mean of differences (B - A) | SD of differences | SEM of differences | 95% confidence interval | R squared (partial eta squared) | Significantly different (P < 0.05)? |
|---|---|---|---|---|---|---|---|---|---|---|---|
| 400 | 0.9333 +/- 1.443 | 1 +/- 1 | 0.9077 | t=0.1311, df=2 | ns | 0.1333 | 1.762 | 1.017 | -4.243 to 4.509 | 0.00852 | No |
| 460 | 1.267 +/- 1.250 | 2.875 +/- 7.411 | 0.2476 | t=1.616, df=2 | ns | 1.833 | 1.966 | 1.135 | -3.049 to 6.716 | 0.5662 | No |
| 480 | 1.800 +/- 0.9539 | 6.225 +/- 1.318 | 0.0486 | t=4.370, df=2 | * | 4.933 | 1.955 | 1.129 | 0.07601 to 9.791 | 0.9052 | Yes |
| 500 | 2.133 +/- 1.124 | 11.38 +/- 1.573 | 0.0202 | t=6.924, df=2 | * | 9.8 | 2.452 | 1.415 | 3.710 to 15.89 | 0.96 | Yes |
| 520 | 3.967 +/- 1.165 | 17.05 +/- 1.923 | 0.0153 | t=7.982, df=2 | * | 13.87 | 3.009 | 1.737 | 6.392 to 21.34 | 0.9696 | Yes |
| 540 | 7.500 +/- 2.166 | 22.13 +/- 3.006 | 0.0359 | t=5.135, df=2 | * | 15.37 | 5.183 | 2.992 | 2.491 to 28.24 | 0.9295 | Yes |
| 560 | 12.03 +/- 2.926 | 28.05 +/-2.745 | 0.0421 | t=4.718, df=2 | * | 15.97 | 5.862 | 3.384 | 1.405 to 30.53 | 0.9175 | Yes |
| 580 | 17.43 +/- 3.502 | 33.13 +/- 2.696 | 0.0434 | t=4.642, df=2 | * | 16.27 | 6.07 | 3.504 | 1.188 to 31.35 | 0.9151 | Yes |
| 600 | 23.20 +/- 3.396 | 34.53 +/- 5.003 | 0.0862 | t=3.182, df=2 | ns | 12.47 | 6.786 | 3.918 | -4.390 to 29.32 | 0.8351 | No |
| 620 | 29.37 +/- 4.222 | 39.33 +/- 4.122 | 0.1092 | t=2.772, df=2 | ns | 11.27 | 7.04 | 4.065 | -6.222 to 28.76 | 0.7935 | No |
| 640 | 38.70 +/- 3.554 | 41.20 +/- 4.925 | 0.4907 | t=0.8370, df=2 | ns | 4 | 8.278 | 4.779 | -16.56 to 24.56 | 0.2594 | No |
| 660 | 43.10 +/- 2.364 | 40.50 +/- 2.576 | 0.2136 | t=1.800, df=2 | ns | -2.667 | 2.566 | 1.481 | -9.040 to 3.707 | 0.6184 | No |
| 680 | 50.37 +/- 2.122 | 42.00 +/- 2.137 | 0.041 | t=4.788, df=2 | * | -7.767 | 2.81 | 1.622 | -14.75 to -0.7875 | 0.9198 | Yes |
| 700 | 56.07 +/- 2.857 | 44.00 +/- 2.423 | 0.0328 | t=5.385, df=2 | * | -11.53 | 3.71 | 2.142 | -20.75 to -2.317 | 0.9355 | Yes |
| 740 | 62.30 +/-3.940 | 48.25 +/- 5.635 | 0.1435 | t=2.347, df=2 | ns | -14.53 | 10.73 | 6.193 | -41.18 to 12.11 | 0.7336 | No |
| 760 | 65.43 +/- 4.839 | 50.25 +/- 3.313 | 0.021 | t=6.795, df=2 | * | -16.77 | 4.274 | 2.467 | -27.38 to -6.151 | 0.9585 | Yes |
| 780 | 67.27 +/- 5.613 | 50.98 +/- 3.015 | 0.0278 | t=5.876, df=2 | * | -17.4 | 5.129 | 2.961 | -30.14 to -4.658 | 0.9452 | Yes |
| 800 | 69.23 +/- 5.719 | 52.78 +/- 3.448 | 0.0461 | t=4.497, df=2 | * | -17.9 | 6.894 | 3.98 | -35.03 to -0.7739 | 0.91 | Yes |
| 820 | 70.17 +/- 6.643 | 53.78 +/- 3.807 | 0.0493 | t=4.336, df=2 | * | -18.13 | 7.243 | 4.182 | -36.13 to -0.1403 | 0.9039 | Yes |
| 830 | 70.6 +/- 6.609 | 59.50 +/- 7.939 | 0.3087 | t=1.353, df=2 | ns | -12.23 | 15.66 | 9.043 | -51.14 to 26.67 | 0.4778 | No |
| 850 | 79.10 +/- 2.152 | 66.10 +/- 3.369 | 0.0128 | t=8.743, df=2 | * | -12.63 | 2.503 | 1.445 | -18.85 to -6.416 | 0.9745 | Yes |

**Supplementary Table 1. Light transmittance measurements between RatDISCO and iDISCO+ cleared tissue with two-tailed paired t-test statistical analysis results.**

Table summarising the two-tailed paired t-test analysis comparing the percentage of light passing through paired brain hemispheres from the same animal cleared with RatDISCO or iDISCO+. We measured wavelengths between 400 and 850 nm at 20-60 nm intervals. The columns indicate the wavelength measured, the mean values and standard deviations (SD) obtained from the spectrophotometer for RatDISCO and iDISCO+cleared tissue; the P values; the t-statistic value and the degrees of freedom t, (df); the P value summary; the mean, SD and standard error (SEM) differences; the 95% confidence interval values, if measurements were significantly different ($P < 0.05$). Columns in orange indicate iDISCO+ measurements with higher light transmittance compared to RatDISCO, while green columns show RatDISCO measurements with greater transmittance than iDISCO+.

| Size in mm | Before clearing | RatDISCO | iDISCO+ |
|---|---|---|---|
| Mean | 22.8 | 17.15 | 20.08 |
| SD | 1.503 | 0.3 | 1.497 |
| SEM | 0.7517 | 0.15 | 0.7487 |
| Lower 95% CI of mean | 20.41 | 16.67 | 17.69 |
| Upper 95% CI of mean | 25.19 | 17.63 | 22.46 |
| Coefficient of variation | 6.59% | 1.75% | 7.46% |
| **ANOVA summary** | | **RatDISCO** | **iDISCO+** |
| F | | 1.095 | 5.731 |
| P value | | 0.3753 | 0.0248 |
| F (DFn, DFd) | | 2.054 (2, 9) | 10.91 (2, 9) |
| P value summary | | ns | * |
| Significant diff. among means (P < 0.05)? | | No | Yes |
| **RatDISCO**<br>**Tukey's multiple comparisons test** | **Mean Diff.** | **95.00% CI of diff.** | **Adjusted P Value** |
| AP vs. ML | 2.65 | -8.210 to 13.51 | 0.7799 |
| AP vs. DV | -3.1 | -13.96 to 7.760 | 0.7141 |
| ML vs. DV | -5.75 | -16.61 to 5.110 | 0.3453 |
| **iDISCO+**<br>**Tukey's multiple comparisons test** | | | |
| AP vs. ML | -8.8 | -16.06 to -1.542 | 0.0199 |
| AP vs. DV | -4.325 | -11.58 to 2.933 | 0.2703 |
| ML vs. DV | 4.475 | -2.783 to 11.73 | 0.2497 |

**Supplementary Table 2. Size measurements of RatDISCO and iDISCO+ cleared rat brain hemispheres with one-way ANOVA analysis results.**

Table summarising the descriptive statistics (green) and the results of one-way ANOVA (blue) comparing brain hemisphere sizes before and after clearing using RatDISCO or iDISCO+. Measurements were taken along the anterior-posterior, mediolateral, and dorsoventral axes for both hemispheres of the same animal. The rows display the mean length (in mm), standard deviation (SD), standard error (SEM), 95% confidence intervals (CI) and the coefficient of variation. The one-way ANOVA (blue) results include F values, P values, degrees of freedom (F (DFn, DFd), and P value summary, indicating significant differences at a threshold of $P < 0.05$. Tukey's multiple comparisons post hoc test results are shown for each pairwise comparison, including mean differences, 95% confidence interval (CI) of the difference, and adjusted P values for RatDISCO and iDISCO+ treated tissue.

| Amygdala | | | | | |
|---|---|---|---|---|---|
| *Source of Variation* | *% of total variation* | *P value* | *P value summary* | *Significant?* | |
| Interaction (genotype vs condition) | 6.5 | 0.1139 | ns | No | |
| Row Factor (WT vs Fmr1-/y ) | 17 | 0.0017 | ** | Yes | |
| Column Factor (naïve, LS, and LS+FS condition) | 46 | <0.0001 | **** | Yes | |
| ***ANOVA table*** | *SS (Type III)* | *DF* | *MS* | *F (DFn, DFd)* | *P value* |
| Interaction | 36 | 2 | 18 | F (2, 24) = 2.4 | P=0.1139 |
| Row Factor | 95 | 1 | 95 | F (1, 24) = 12 | P=0.0017 |
| Column Factor | 256 | 2 | 128 | F (2, 24) = 17 | P<0.0001 |
| Residual | 182 | 24 | 7.6 | | |
| ***Tukey's multiple comparisons test*** | *Predicted (LS) mean diff.* | *95.00% CI of diff.* | *Below threshold?* | *Summary* | *Adjusted P Value* |
| WT:Naive vs. WT:LS | -5.2 | -10 to -0.054 | Yes | * | 0.0467 |
| WT:Naive vs. WT:LS+FS | -7.1 | -13 to -1.6 | Yes | ** | 0.006 |
| WT:Naive vs. Fmr1-/y :Naive | 2.1 | -3.1 to 7.3 | No | ns | 0.8025 |
| WT:Naive vs. Fmr1-/y:LS | 1.5 | -3.7 to 6.6 | No | ns | 0.946 |
| WT:Naive vs. Fmr1-/y:LS+FS | -5.2 | -10 to -0.040 | Yes | * | 0.0475 |
| WT:LS vs. WT:LS+FS | -1.9 | -7.6 to 3.8 | No | ns | 0.8997 |
| WT:LS vs.Fmr1-/y:Naive | 7.3 | 1.9 to 13 | Yes | ** | 0.0038 |
| WT:LS vs. Fmr1-/y:LS | 6.7 | 1.3 to 12 | Yes | ** | 0.0091 |
| WT:LS vs. Fmr1-/y:LS+FS | 0.014 | -5.4 to 5.4 | No | ns | >0.9999 |
| WT:LS+FS vs. Fmr1-/y:Naive | 9.2 | 3.5 to 15 | Yes | *** | 0.0005 |
| WT:LS+FS vs. Fmr1-/y:LS | 8.6 | 2.9 to 14 | Yes | ** | 0.0012 |
| WT:LS+FS vs.Fmr1-/y:LS+FS | 1.9 | -3.8 to 7.6 | No | ns | 0.8969 |
| Fmr1-/y:Naive vs. Fmr1-/y:LS | -0.62 | -6.0 to 4.8 | No | ns | 0.9991 |
| Fmr1-/y:Naive vs. Fmr1-/y:LS+FS | -7.3 | -13 to -1.9 | Yes | ** | 0.0039 |
| Fmr1-/y:LS vs. Fmr1-/y:LS+FS | -6.7 | -12 to -1.3 | Yes | ** | 0.0093 |

**Supplementary Table 3. Two-Way ANOVA analysis of cFOS cell density in the amygdala of WT and *Fmr1$^{-/y}$* rats.**

Summary table of 2-way ANOVA results and Tukey's multiple comparisons test. cFOS cell density was measured under naïve (no stimulus), exposure to flashing light stimuli (LS), and exposure to LS combined with a footshock (LS+FS) experimental conditions. 2-way ANOVA (alpha 0.05) results show the percentage of variation (%), P-value, P value summary and significant differences at a threshold of $P < 0.05$ per interaction, the sum of squares (SS), mean squares (MS), degrees of freedom (DS), F (DFn, DFd), and P value of the interaction between genotype vs condition, the genotype and the experimental condition effects. Tukey's multiple comparisons post hoc test results are shown for each pairwise comparison, including predicted mean differences, 95% CI of the difference, threshold and summary, and adjusted P-values.

| Basal complex | | | | | |
|---|---|---|---|---|---|
| *Source of Variation* | *% of total variation* | *P value* | *P value summary* | *Significant?* | |
| Interaction | 7.334 | 0.116 | ns | No | |
| Genotype | 15.82 | 0.0039 | ** | Yes | |
| Treatment | 40.7 | 0.0001 | *** | Yes | |
| **ANOVA table** | SS (Type III) | DF | MS | F (DFn, DFd) | P value |
| Interaction | 28.13 | 2 | 14.06 | F (2, 24) = 2.360 | P=0.1160 |
| Genotype | 60.68 | 1 | 60.68 | F (1, 24) = 10.18 | P=0.0039 |
| Treatment | 156.1 | 2 | 78.04 | F (2, 24) = 13.10 | P=0.0001 |
| Residual | 143 | 24 | 5.96 | | |
| ***Tukey's multiple comparisons test*** | *Predicted (LS) mean diff.* | *95.00% CI of diff.* | *Below threshold?* | *Summary* | *Adjusted P Value* |
| WT:Naive vs. WT:LS | -3.5 | -8.1 to 1.0 | No | ns | 0.1987 |
| WT:Naive vs. WT:LS+FS | -5.3 | -10 to -0.38 | Yes | * | 0.0294 |
| WT:Naive vs. Fmr1-/y :Naive | 1.8 | -2.8 to 6.4 | No | ns | 0.8253 |
| WT:Naive vs. Fmr1-/y:LS | 2.1 | -2.5 to 6.6 | No | ns | 0.7263 |
| WT:Naive vs. Fmr1-/y:LS+FS | -4.1 | -8.6 to 0.51 | No | ns | 0.102 |
| WT:LS vs. WT:LS+FS | -1.7 | -6.8 to 3.3 | No | ns | 0.8967 |
| WT:LS vs.Fmr1-/y:Naive | 5.3 | 0.56 to 10 | Yes | * | 0.0224 |
| WT:LS vs. Fmr1-/y:LS | 5.6 | 0.83 to 10 | Yes | * | 0.0149 |
| WT:LS vs. Fmr1-/y:LS+FS | -0.53 | -5.3 to 4.2 | No | ns | 0.9993 |
| WT:LS+FS vs. Fmr1-/y:Naive | 7 | 2.0 to 12 | Yes | ** | 0.003 |
| WT:LS+FS vs. Fmr1-/y:LS | 7.3 | 2.3 to 12 | Yes | ** | 0.002 |
| WT:LS+FS vs.Fmr1-/y:LS+FS | 1.2 | -3.9 to 6.3 | No | ns | 0.9766 |
| Fmr1-/y:Naive vs. Fmr1-/y:LS | 0.27 | -4.5 to 5.0 | No | ns | >0.9999 |
| Fmr1-/y:Naive vs. Fmr1-/y:LS+FS | -5.9 | -11 to -1.1 | Yes | * | 0.0101 |
| Fmr1-/y:LS vs. Fmr1-/y:LS+FS | -6.1 | -11 to -1.4 | Yes | ** | 0.0066 |

**Supplementary Table 4. Two-way ANOVA analysis of cFOS cell density quantification in the basal complex of the amygdala.**

Summary table of 2-way ANOVA results and Tukey's multiple comparisons test. cFOS cell density was measured under naïve (no stimulus), exposure to flashing light stimuli (LS), and exposure to LS combined with a footshock (LS+FS) experimental conditions. 2-way ANOVA (alpha 0.05) results show the percentage of variation (%), P-value, P value summary and significant differences at a threshold of P < 0.05 per interaction, the sum of squares (SS), mean squares (MS), degrees of freedom (DS), F (DFn, DFd), and P value of the interaction between genotype vs condition, the genotype and the experimental condition effects. Tukey's multiple comparisons post hoc test results are shown for each pairwise comparison, including predicted mean differences, 95% CI of the difference, threshold and summary, and adjusted P-values.

| Basomedial complex | | | | | |
|---|---|---|---|---|---|
| Source of Variation | % of total variation | P value | P value summary | Significant? | |
| Interaction | 5.6 | 0.2028 | ns | No | |
| Genotype | 14 | 0.0077 | ** | Yes | |
| Condition (naive, LS or LS+FS) | 43 | 0.0001 | *** | Yes | |
| **ANOVA table** | SS (Type III) | DF | MS | F (DFn, DFd) | P value |
| Interaction | 62 | 2 | 31 | F (2, 24) = 1.7 | P=0.2028 |
| Genotype | 155 | 1 | 155 | F (1, 24) = 8.5 | P=0.0077 |
| Condition (naive, LS or LS+FS) | 482 | 2 | 241 | F (2, 24) = 13 | P=0.0001 |
| Residual | 439 | 24 | 18 | | |
| **Tukey's multiple comparisons test** | Predicted (LS) mean diff. | 95.00% CI of diff. | Below threshold? | Summary | Adjusted P Value |
| WT:Naive vs. WT:CS | -8.2 | -16 to -0.16 | Yes | * | 0.0438 |
| WT:Naive vs. WT:CS+US | -10 | -19 to -1.5 | Yes | * | 0.0142 |
| WT:Naive vs. Fmr1 KO:Naive | 2.4 | -5.7 to 10 | No | ns | 0.9402 |
| WT:Naive vs. Fmr1 KO:CS | 0.48 | -7.5 to 8.5 | No | ns | >0.9999 |
| WT:Naive vs. Fmr1 KO:CS+US | -7.4 | -15 to 0.65 | No | ns | 0.0844 |
| WT:CS vs. WT:CS+US | -1.9 | -11 to 7.0 | No | ns | 0.9839 |
| WT:CS vs. Fmr1 KO:Naive | 11 | 2.2 to 19 | Yes | ** | 0.0081 |
| WT:CS vs. Fmr1 KO:CS | 8.6 | 0.28 to 17 | Yes | * | 0.04 |
| WT:CS vs. Fmr1 KO:CS+US | 0.81 | -7.6 to 9.2 | No | ns | 0.9996 |
| WT:CS+US vs. Fmr1 KO:Naive | 12 | 3.6 to 21 | Yes | ** | 0.0028 |
| WT:CS+US vs. Fmr1 KO:CS | 11 | 1.7 to 19 | Yes | * | 0.0133 |
| WT:CS+US vs. Fmr1 KO:CS+US | 2.7 | -6.2 to 12 | No | ns | 0.9296 |
| Fmr1 KO:Naive vs. Fmr1 KO:CS | -1.9 | -10 to 6.5 | No | ns | 0.9808 |
| Fmr1 KO:Naive vs. Fmr1 KO:CS+US | -9.7 | -18 to -1.4 | Yes | * | 0.0163 |
| Fmr1 KO:CS vs. Fmr1 KO:CS+US | -7.8 | -16 to 0.53 | No | ns | 0.0754 |

**Supplementary Table 5. Two-way ANOVA analysis of cFOS cell density quantification in the basomedial complex of the amygdala.**

Summary table of 2-way ANOVA results and Tukey's multiple comparisons test. cFOS cell density was measured under naïve (no stimulus), exposure to flashing light stimuli (LS), and exposure to LS combined with a footshock (LS+FS) experimental conditions. 2-way ANOVA (alpha 0.05) results show the percentage of variation (%), P-value, P value summary and significant differences at a threshold of $P < 0.05$ per interaction, the sum of squares (SS), mean squares (MS), degrees of freedom (DS), F (DFn, DFd), and P value of the interaction between genotype vs condition, the genotype and the experimental condition effects. Tukey's multiple comparisons post hoc test results are shown for each pairwise comparison, including predicted mean differences, 95% CI of the difference, threshold and summary, and adjusted P-values.

| Central complex | | | | | |
|---|---|---|---|---|---|
| *Source of Variation* | *% of total variation* | *P value* | *P value summary* | *Significant?* | |
| Interaction | 3.1 | 0.603 | ns | No | |
| Genotype | 2.3 | 0.3858 | ns | No | |
| Treatment | 22 | 0.0399 | * | Yes | |
| ANOVA table | SS (Type III) | DF | MS | F (DFn, DFd) | P value |
| Interaction | 6.2 | 2 | 3.1 | F (2, 24) = 0.52 | P=0.6030 |
| Genotype | 4.7 | 1 | 4.7 | F (1, 24) = 0.78 | P=0.3858 |
| Treatment | 45 | 2 | 22 | F (2, 24) = 3.7 | P=0.0399 |
| Residual | 145 | 24 | 6 | | |

| Lateral complex | | | | | |
|---|---|---|---|---|---|
| *Source of Variation* | *% of total variation* | *P value* | *P value summary* | *Significant?* | |
| Interaction | 0.78 | 0.8772 | ns | No | |
| Genotype | 19 | 0.0186 | * | Yes | |
| Treatment | 10 | 0.1933 | ns | No | |
| ANOVA table | SS (Type III) | DF | MS | F (DFn, DFd) | P value |
| Interaction | 2.7 | 2 | 1.4 | F (2, 24) = 0.13 | P=0.8772 |
| Genotype | 66 | 1 | 66 | F (1, 24) = 6.4 | P=0.0186 |
| Treatment | 36 | 2 | 18 | F (2, 24) = 1.8 | P=0.1933 |
| Residual | 248 | 24 | 10 | | |

**Supplementary Table 6. Two-way ANOVA analysis of cFOS cell density quantification in the central and lateral complex of the amygdala.**

Summary tables of 2-way ANOVA results. cFOS cell density was measured under naïve (no stimulus), exposure to flashing light stimuli (LS), and exposure to LS combined with a footshock (LS+FS) experimental conditions. 2-way ANOVA (alpha 0.05) results show the percentage of variation (%), P-value, P value summary and significant differences at a threshold of $P < 0.05$ per interaction, the sum of squares (SS), mean squares (MS), degrees of freedom (DS), F (DFn, DFd), and P value of the interaction between genotype vs condition, the genotype and the experimental condition effects. Tukey's multiple comparisons post hoc test results showed no significant differences for any pairwise comparison (data not shown).

| *BLA subnuclei* | | | | | |
|---|---|---|---|---|---|
| *Source of Variation* | *% of total variation* | *P value* | *P value summary* | *Significant?* | |
| Interaction | 4.8 | 0.2643 | ns | No | |
| Genotype | 7.9 | 0.0425 | * | Yes | |
| Treatment | 46 | 0.0001 | *** | Yes | |
| ANOVA table | SS (Type III) | DF | MS | F (DFn, DFd) | P value |
| Interaction | 15 | 2 | 7.3 | F (2, 24) = 1.4 | P=0.2643 |
| Genotype | 24 | 1 | 24 | F (1, 24) = 4.6 | P=0.0425 |
| Treatment | 140 | 2 | 70 | F (2, 24) = 13 | P=0.0001 |
| Residual | 125 | 24 | 5.2 | | |
| *Tukey's multiple comparisons test* | *Predicted (LS) mean diff.* | *95.00% CI of diff.* | *Below threshold?* | *Summary* | *Adjusted P Value* |
| WT:Naive vs. WT:LS | -1.4 | -5.6 to 2.9 | No | ns | 0.9204 |
| WT:Naive vs. WT:LS+FS | -4.5 | -9.0 to 0.10 | No | ns | 0.0581 |
| WT:Naive vs. Fmr1-/y :Naive | 1.1 | -3.2 to 5.4 | No | ns | 0.9649 |
| WT:Naive vs. Fmr1-/y:LS | 2.4 | -1.9 to 6.7 | No | ns | 0.5145 |
| WT:Naive vs. Fmr1-/y:LS+FS | -3.9 | -8.2 to 0.34 | No | ns | 0.0838 |
| WT:LS vs. WT:LS+FS | -3.1 | -7.8 to 1.6 | No | ns | 0.3583 |
| WT:LS vs.Fmr1-/y:Naive | 2.5 | -2.0 to 6.9 | No | ns | 0.5436 |
| WT:LS vs. Fmr1-/y:LS | 3.8 | -0.69 to 8.2 | No | ns | 0.1329 |
| WT:LS vs. Fmr1-/y:LS+FS | -2.6 | -7.0 to 1.9 | No | ns | 0.4916 |
| WT:LS+FS vs. Fmr1-/y:Naive | 5.6 | 0.82 to 10 | Yes | * | 0.0149 |
| WT:LS+FS vs. Fmr1-/y:LS | 6.9 | 2.1 to 12 | Yes | ** | 0.0019 |
| WT:LS+FS vs.Fmr1-/y:LS+FS | 0.52 | -4.2 to 5.3 | No | ns | 0.9993 |
| Fmr1-/y:Naive vs. Fmr1-/y:LS | 1.3 | -3.2 to 5.8 | No | ns | 0.9402 |
| Fmr1-/y:Naive vs. Fmr1-/y:LS+FS | -5 | -9.5 to -0.57 | Yes | * | 0.0206 |
| Fmr1-/y:LS vs. Fmr1-/y:LS+FS | -6.4 | -11 to -1.9 | Yes | ** | 0.0023 |

**Supplementary Table 7. Two-way ANOVA analysis of cFOS cell density quantification in the BLA subnuclei of the amygdala.**

Summary table of 2-way ANOVA results and Tukey's multiple comparisons test. cFOS cell density was measured under naïve (no stimulus), exposure to flashing light stimuli (LS), and exposure to LS combined with a footshock (LS+FS) experimental conditions. 2-way ANOVA (alpha 0.05) results show the percentage of variation (%), P-value, P value summary and significant differences at a threshold of $P < 0.05$ per interaction, the sum of squares (SS), mean squares (MS), degrees of freedom (DS), F (DFn, DFd), and P value of the interaction between genotype vs condition, the genotype and the experimental condition effects. Tukey's multiple comparisons post hoc test results are shown for each pairwise comparison, including predicted mean differences, 95% CI of the difference, threshold and summary, and adjusted P-values.

| BLV subnuclei | | | | | |
|---|---|---|---|---|---|
| *Source of Variation* | *% of total variation* | *P value* | *P value summary* | *Significant?* | |
| Interaction | 10 | 0.0209 | * | Yes | |
| Genotype | 2 | 0.2004 | ns | No | |
| Treatment | 58 | <0.0001 | **** | Yes | |
| ANOVA table | SS (Type III) | DF | MS | F (DFn, DFd) | P value |
| Interaction | 171 | 2 | 86 | F (2, 24) = 4.6 | P=0.0209 |
| Genotype | 33 | 1 | 33 | F (1, 24) = 1.7 | P=0.2004 |
| Treatment | 946 | 2 | 473 | F (2, 24) = 25 | P<0.0001 |
| Residual | 450 | 24 | 19 | | |
| *Tukey's multiple comparisons test* | *Predicted (LS) mean diff.* | *95.00% CI of diff.* | *Below threshold?* | *Summary* | *Adjusted P Value* |
| WT:Naive vs. WT:LS | -7.1 | -15 to 1.0 | No | ns | 0.1129 |
| WT:Naive vs. WT:LS+FS | -10 | -19 to -1.4 | Yes | * | 0.0161 |
| WT:Naive vs. Fmr1-/y :Naive | 3.1 | -5.0 to 11 | No | ns | 0.8395 |
| WT:Naive vs. Fmr1-/y:LS | 0.51 | -7.6 to 8.6 | No | ns | >0.9999 |
| WT:Naive vs. Fmr1-/y:LS+FS | -14 | -23 to -6.3 | Yes | *** | 0.0002 |
| WT:LS vs. WT:LS+FS | -3 | -12 to 6.0 | No | ns | 0.9042 |
| WT:LS vs.Fmr1-/y:Naive | 10 | 1.7 to 19 | Yes | * | 0.0122 |
| WT:LS vs. Fmr1-/y:LS | 7.6 | -0.89 to 16 | No | ns | 0.0979 |
| WT:LS vs. Fmr1-/y:LS+FS | -7.4 | -16 to 1.1 | No | ns | 0.1129 |
| WT:LS+FS vs. Fmr1-/y:Naive | 13 | 4.2 to 22 | Yes | ** | 0.0017 |
| WT:LS+FS vs. Fmr1-/y:LS | 11 | 1.6 to 20 | Yes | * | 0.0147 |
| WT:LS+FS vs.Fmr1-/y:LS+FS | -4.4 | -13 to 4.6 | No | ns | 0.6584 |
| Fmr1-/y:Naive vs. Fmr1-/y:LS | -2.6 | -11 to 5.9 | No | ns | 0.9296 |
| Fmr1-/y:Naive vs. Fmr1-/y:LS+FS | -18 | -26 to -9.1 | Yes | **** | <0.0001 |
| Fmr1-/y:LS vs. Fmr1-/y:LS+FS | -15 | -23 to -6.5 | Yes | *** | 0.0002 |

**Supplementary Table 8. Two-way ANOVA analysis of cFOS cell density quantification in the BLV subnuclei of the amygdala.**

Summary table of 2-way ANOVA results and Tukey’s multiple comparisons test. cFOS cell density was measured under naïve (no stimulus), exposure to flashing light stimuli (LS), and exposure to LS combined with a footshock (LS+FS) experimental conditions. 2-way ANOVA (alpha 0.05) results show the percentage of variation (%), P-value, P value summary and significant differences at a threshold of P < 0.05 per interaction, the sum of squares (SS), mean squares (MS), degrees of freedom (DS), F (DFn, DFd), and P value of the interaction between genotype vs condition, the genotype and the experimental condition effects. Tukey's multiple comparisons post hoc test results are shown for each pairwise comparison, including predicted mean differences, 95% CI of the difference, threshold and summary, and adjusted P-values.

| BLP subnuclei | | | | | |
|---|---|---|---|---|---|
| *Source of Variation* | *% of total variation* | *P value* | *P value summary* | *Significant?* | |
| Interaction | 6.3 | 0.2632 | ns | No | |
| Genotype | 29 | 0.0014 | ** | Yes | |
| Treatment | 13 | 0.0778 | ns | No | |
| ANOVA table | SS (Type III) | DF | MS | F (DFn, DFd) | P value |
| Interaction | 48 | 2 | 24 | F (2, 24) = 1.4 | P=0.2632 |
| Genotype | 224 | 1 | 224 | F (1, 24) = 13 | P=0.0014 |
| Treatment | 97 | 2 | 49 | F (2, 24) = 2.8 | P=0.0778 |
| Residual | 409 | 24 | 17 | | |
| *Tukey's multiple comparisons test* | *Predicted (LS) mean diff.* | *95.00% CI of diff.* | *Below threshold?* | *Summary* | *Adjusted P Value* |
| WT:Naive vs. WT:LS | -6.8 | -15 to 0.95 | No | ns | 0.1094 |
| WT:Naive vs. WT:LS+FS | -4.9 | -13 to 3.4 | No | ns | 0.4723 |
| WT:Naive vs. Fmr1-/y :Naive | 2.7 | -5.0 to 10 | No | ns | 0.8782 |
| WT:Naive vs. Fmr1-/y:LS | 2 | -5.7 to 9.7 | No | ns | 0.9652 |
| WT:Naive vs. Fmr1-/y:LS+FS | 0.14 | -7.6 to 7.9 | No | ns | >0.9999 |
| WT:LS vs. WT:LS+FS | 1.9 | -6.6 to 10 | No | ns | 0.9806 |
| WT:LS vs.Fmr1-/y:Naive | 9.5 | 1.4 to 18 | Yes | * | 0.0143 |
| WT:LS vs. Fmr1-/y:LS | 8.8 | 0.70 to 17 | Yes | * | 0.0277 |
| WT:LS vs. Fmr1-/y:LS+FS | 6.9 | -1.2 to 15 | No | ns | 0.1234 |
| WT:LS+FS vs. Fmr1-/y:Naive | 7.6 | -0.97 to 16 | No | ns | 0.1034 |
| WT:LS+FS vs. Fmr1-/y:LS | 6.8 | -1.7 to 15 | No | ns | 0.1724 |
| WT:LS+FS vs.Fmr1-/y:LS+FS | 5 | -3.6 to 14 | No | ns | 0.4831 |
| Fmr1-/y:Naive vs. Fmr1-/y:LS | -0.75 | -8.8 to 7.3 | No | ns | 0.9997 |
| Fmr1-/y:Naive vs. Fmr1-/y:LS+FS | -2.6 | -11 to 5.5 | No | ns | 0.9148 |
| Fmr1-/y:LS vs. Fmr1-/y:LS+FS | -1.9 | -9.9 to 6.2 | No | ns | 0.9789 |

**Supplementary Table 9. Two-way ANOVA analysis of cFOS cell density quantification in the BLP subnuclei of the amygdala.**

Summary table of 2-way ANOVA results and Tukey’s multiple comparisons test. cFOS cell density was measured under naïve (no stimulus), exposure to flashing light stimuli (LS), and exposure to LS combined with a footshock (LS+FS) experimental conditions. 2-way ANOVA (alpha 0.05) results show the percentage of variation (%), P-value, P value summary and significant differences at a threshold of P < 0.05 per interaction, the sum of squares (SS), mean squares (MS), degrees of freedom (DS), F (DFn, DFd), and P value of the interaction between genotype vs condition, the genotype and the experimental condition effects. Tukey's multiple comparisons post hoc test results are shown for each pairwise comparison, including predicted mean differences, 95% CI of the difference, threshold and summary, and adjusted P-values.

| BMA subnuclei | | | | | |
|---|---|---|---|---|---|
| *Source of Variation* | *% of total variation* | *P value* | *P value summary* | *Significant?* | |
| Interaction | 6.7 | 0.2096 | ns | No | |
| Genotype | 7.3 | 0.068 | ns | No | |
| Treatment | 39 | 0.0009 | *** | Yes | |
| ANOVA table | SS (Type III) | DF | MS | F (DFn, DFd) | P value |
| Interaction | 104 | 2 | 52 | F (2, 24) = 1.7 | P=0.2096 |
| Genotype | 114 | 1 | 114 | F (1, 24) = 3.7 | P=0.0680 |
| Treatment | 598 | 2 | 299 | F (2, 24) = 9.6 | P=0.0009 |
| Residual | 747 | 24 | 31 | | |
| *Tukey's multiple comparisons test* | *Predicted (LS) mean diff.* | *95.00% CI of diff.* | *Below threshold?* | *Summary* | *Adjusted P Value* |
| WT:Naive vs. WT:LS | -9.8 | -20 to 0.62 | No | ns | 0.0735 |
| WT:Naive vs. WT:LS+FS | -12 | -23 to -0.76 | Yes | * | 0.0314 |
| WT:Naive vs. Fmr1-/y :Naive | 0.5 | -9.9 to 11 | No | ns | >0.9999 |
| WT:Naive vs. Fmr1-/y:LS | -0.79 | -11 to 9.7 | No | ns | 0.9999 |
| WT:Naive vs. Fmr1-/y:LS+FS | -9.7 | -20 to 0.77 | No | ns | 0.0809 |
| WT:LS vs. WT:LS+FS | -2.1 | -14 to 9.5 | No | ns | 0.9931 |
| WT:LS vs.Fmr1-/y:Naive | 10 | -0.59 to 21 | No | ns | 0.071 |
| WT:LS vs. Fmr1-/y:LS | 9 | -1.9 to 20 | No | ns | 0.1459 |
| WT:LS vs. Fmr1-/y:LS+FS | 0.16 | -11 to 11 | No | ns | >0.9999 |
| WT:LS+FS vs. Fmr1-/y:Naive | 12 | 0.82 to 24 | Yes | * | 0.0309 |
| WT:LS+FS vs. Fmr1-/y:LS | 11 | -0.46 to 23 | No | ns | 0.065 |
| WT:LS+FS vs.Fmr1-/y:LS+FS | 2.2 | -9.3 to 14 | No | ns | 0.9904 |
| Fmr1-/y:Naive vs. Fmr1-/y:LS | -1.3 | -12 to 9.6 | No | ns | 0.9991 |
| Fmr1-/y:Naive vs. Fmr1-/y:LS+FS | -10 | -21 to 0.74 | No | ns | 0.0778 |
| Fmr1-/y:LS vs. Fmr1-/y:LS+FS | -8.9 | -20 to 2.0 | No | ns | 0.1584 |

**Supplementary Table 10. Two-way ANOVA analysis of cFOS cell density quantification in the BMA subnuclei of the amygdala.**

Summary table of 2-way ANOVA results and Tukey's multiple comparisons test. cFOS cell density was measured under naïve (no stimulus), exposure to flashing light stimuli (LS), and exposure to LS combined with a footshock (LS+FS) experimental conditions. 2-way ANOVA (alpha 0.05) results show the percentage of variation (%), P-value, P value summary and significant differences at a threshold of $P < 0.05$ per interaction, the sum of squares (SS), mean squares (MS), degrees of freedom (DS), F (DFn, DFd), and P value of the interaction between genotype vs condition, the genotype and the experimental condition effects. Tukey's multiple comparisons post hoc test results are shown for each pairwise comparison, including predicted mean differences, 95% CI of the difference, threshold and summary, and adjusted P-values.

| BMP subnuclei | | | | | |
|---|---|---|---|---|---|
| *Source of Variation* | *% of total variation* | *P value* | *P value summary* | *Significant?* | |
| Interaction | 2.7 | 0.52 | ns | No | |
| Genotype | 20 | 0.0047 | ** | Yes | |
| Treatment | 32 | 0.0023 | ** | Yes | |
| ANOVA table | SS (Type III) | DF | MS | F (DFn, DFd) | P value |
| Interaction | 30 | 2 | 15 | F (2, 24) = 0.67 | P=0.5200 |
| Genotype | 216 | 1 | 216 | F (1, 24) = 9.7 | P=0.0047 |
| Treatment | 352 | 2 | 176 | F (2, 24) = 7.9 | P=0.0023 |
| Residual | 535 | 24 | 22 | | |
| *Tukey's multiple comparisons test* | *Predicted (LS) mean diff.* | *95.00% CI of diff.* | *Below threshold?* | *Summary* | *Adjusted P Value* |
| WT:Naive vs. WT:LS | -6.2 | -15 to 2.7 | No | ns | 0.292 |
| WT:Naive vs. WT:LS+FS | -7.9 | -17 to 1.5 | No | ns | 0.1389 |
| WT:Naive vs. Fmr1-/y :Naive | 4.6 | -4.3 to 13 | No | ns | 0.6046 |
| WT:Naive vs. Fmr1-/y:LS | 2 | -6.8 to 11 | No | ns | 0.979 |
| WT:Naive vs. Fmr1-/y:LS+FS | -4.4 | -13 to 4.4 | No | ns | 0.6343 |
| WT:LS vs. WT:LS+FS | -1.7 | -12 to 8.1 | No | ns | 0.9938 |
| WT:LS vs.Fmr1-/y:Naive | 11 | 1.5 to 20 | Yes | * | 0.0159 |
| WT:LS vs. Fmr1-/y:LS | 8.2 | -1.0 to 17 | No | ns | 0.1021 |
| WT:LS vs. Fmr1-/y:LS+FS | 1.7 | -7.5 to 11 | No | ns | 0.9914 |
| WT:LS+FS vs. Fmr1-/y:Naive | 12 | 2.7 to 22 | Yes | ** | 0.0072 |
| WT:LS+FS vs. Fmr1-/y:LS | 9.9 | 0.12 to 20 | Yes | * | 0.046 |
| WT:LS+FS vs.Fmr1-/y:LS+FS | 3.4 | -6.3 to 13 | No | ns | 0.8816 |
| Fmr1-/y:Naive vs. Fmr1-/y:LS | -2.6 | -12 to 6.7 | No | ns | 0.9534 |
| Fmr1-/y:Naive vs. Fmr1-/y:LS+FS | -9 | -18 to 0.21 | No | ns | 0.058 |
| Fmr1-/y:LS vs. Fmr1-/y:LS+FS | -6.5 | -16 to 2.8 | No | ns | 0.2887 |

**Supplementary Table 11. Two-way ANOVA analysis of cFOS cell density quantification in the BMP subnuclei of the amygdala.**

Summary table of 2-way ANOVA results and Tukey's multiple comparisons test. cFOS cell density was measured under naïve (no stimulus), exposure to flashing light stimuli (LS), and exposure to LS combined with a footshock (LS+FS) experimental conditions. 2-way ANOVA (alpha 0.05) results show the percentage of variation (%), P-value, P value summary and significant differences at a threshold of P < 0.05 per interaction, the sum of squares (SS), mean squares (MS), degrees of freedom (DS), F (DFn, DFd), and P value of the interaction between genotype vs condition, the genotype and the experimental condition effects. Tukey's multiple comparisons post hoc test results are shown for each pairwise comparison, including predicted mean differences, 95% CI of the difference, threshold and summary, and adjusted P-values.

| LaDL subnuclei | | | | | |
|---|---|---|---|---|---|
| *Source of Variation* | *% of total variation* | *P value* | *P value summary* | *Significant?* | |
| Interaction | 2.8 | 0.6481 | ns | No | |
| Genotype | 14 | 0.0488 | * | Yes | |
| Treatment | 8.4 | 0.2836 | ns | No | |
| ANOVA table | SS (Type III) | DF | MS | F (DFn, DFd) | P value |
| Interaction | 9.7 | 2 | 4.8 | F (2, 24) = 0.44 | P=0.6481 |
| Genotype | 47 | 1 | 47 | F (1, 24) = 4.3 | P=0.0488 |
| Treatment | 29 | 2 | 15 | F (2, 24) = 1.3 | P=0.2836 |
| Residual | 262 | 24 | 11 | | |
| *Tukey's multiple comparisons test* | *Predicted (LS) mean diff.* | *95.00% CI of diff.* | *Below threshold?* | *Summary* | *Adjusted P Value* |
| WT:Naive vs. WT:LS | 2.6 | -3.6 to 8.8 | No | ns | 0.78 |
| WT:Naive vs. WT:LS+FS | -1.2 | -7.8 to 5.4 | No | ns | 0.99 |
| WT:Naive vs. Fmr1-/y :Naive | 2.9 | -3.2 to 9.1 | No | ns | 0.69 |
| WT:Naive vs. Fmr1-/y:LS | 3.6 | -2.6 to 9.7 | No | ns | 0.50 |
| WT:Naive vs. Fmr1-/y:LS+FS | 2.5 | -3.7 to 8.7 | No | ns | 0.81 |
| WT:LS vs. WT:LS+FS | -3.8 | -11 to 3.1 | No | ns | 0.54 |
| WT:LS vs.Fmr1-/y:Naive | 0.34 | -6.1 to 6.8 | No | ns | >0.9999 |
| WT:LS vs. Fmr1-/y:LS | 0.95 | -5.5 to 7.4 | No | ns | 1.00 |
| WT:LS vs. Fmr1-/y:LS+FS | -0.1 | -6.6 to 6.4 | No | ns | >0.9999 |
| WT:LS+FS vs. Fmr1-/y:Naive | 4.1 | -2.7 to 11 | No | ns | 0.45 |
| WT:LS+FS vs. Fmr1-/y:LS | 4.7 | -2.1 to 12 | No | ns | 0.30 |
| WT:LS+FS vs.Fmr1-/y:LS+FS | 3.7 | -3.2 to 11 | No | ns | 0.57 |
| Fmr1-/y:Naive vs. Fmr1-/y:LS | 0.61 | -5.9 to 7.1 | No | ns | 1.00 |
| Fmr1-/y:Naive vs. Fmr1-/y:LS+FS | -0.44 | -6.9 to 6.0 | No | ns | >0.9999 |
| Fmr1-/y:LS vs. Fmr1-/y:LS+FS | -1.1 | -7.5 to 5.4 | No | ns | 1.00 |

**Supplementary Table 12. Two-way ANOVA analysis of cFOS cell density quantification in the LaDL subnuclei of the amygdala.**

Summary table of 2-way ANOVA results and Tukey's multiple comparisons test. cFOS cell density was measured under naïve (no stimulus), exposure to flashing light stimuli (LS), and exposure to LS combined with a footshock (LS+FS) experimental conditions. 2-way ANOVA (alpha 0.05) results show the percentage of variation (%), P-value, P value summary and significant differences at a threshold of P < 0.05 per interaction, the sum of squares (SS), mean squares (MS), degrees of freedom (DS), F (DFn, DFd), and P value of the interaction between genotype vs condition, the genotype and the experimental condition effects. Tukey's multiple comparisons post hoc test results are shown for each pairwise comparison, including predicted mean differences, 95% CI of the difference, threshold and summary, and adjusted P-values.

| *LaVM subnuclei* | | | | | |
|---|---|---|---|---|---|
| *Source of Variation* | *% of total variation* | *P value* | *P value summary* | *Significant?* | |
| Interaction | 0.38 | 0.9357 | ns | No | |
| Genotype | 21 | 0.0116 | * | Yes | |
| Treatment | 11 | 0.1685 | ns | No | |
| ANOVA table | SS (Type III) | DF | MS | F (DFn, DFd) | P value |
| Interaction | 1.6 | 2 | 0.78 | F (2, 24) = 0.067 | P=0.9357 |
| Genotype | 88 | 1 | 88 | F (1, 24) = 7.5 | P=0.0116 |
| Treatment | 45 | 2 | 23 | F (2, 24) = 1.9 | P=0.1685 |
| Residual | 282 | 24 | 12 | | |
| *Tukey's multiple comparisons test* | *Predicted (LS) mean diff.* | *95.00% CI of diff.* | *Below threshold?* | *Summary* | *Adjusted P Value* |
| WT:Naive vs. WT:LS | 1 | -5.4 to 7.5 | No | ns | 1.00 |
| WT:Naive vs. WT:LS+FS | -2.2 | -9.1 to 4.6 | No | ns | 0.92 |
| WT:Naive vs. Fmr1-/y :Naive | 2.8 | -3.6 to 9.2 | No | ns | 0.75 |
| WT:Naive vs. Fmr1-/y:LS | 4.7 | -1.8 to 11 | No | ns | 0.26 |
| WT:Naive vs. Fmr1-/y:LS+FS | 1.7 | -4.7 to 8.1 | No | ns | 0.96 |
| WT:LS vs. WT:LS+FS | -3.2 | -10 to 3.9 | No | ns | 0.72 |
| WT:LS vs.Fmr1-/y:Naive | 1.8 | -4.9 to 8.5 | No | ns | 0.96 |
| WT:LS vs. Fmr1-/y:LS | 3.6 | -3.1 to 10 | No | ns | 0.56 |
| WT:LS vs. Fmr1-/y:LS+FS | 0.66 | -6.1 to 7.4 | No | ns | 1.00 |
| WT:LS+FS vs. Fmr1-/y:Naive | 5 | -2.1 to 12 | No | ns | 0.28 |
| WT:LS+FS vs. Fmr1-/y:LS | 6.9 | -0.26 to 14 | No | ns | 0.06 |
| WT:LS+FS vs.Fmr1-/y:LS+FS | 3.9 | -3.2 to 11 | No | ns | 0.55 |
| Fmr1-/y:Naive vs. Fmr1-/y:LS | 1.8 | -4.9 to 8.5 | No | ns | 0.96 |
| Fmr1-/y:Naive vs. Fmr1-/y:LS+FS | -1.1 | -7.8 to 5.6 | No | ns | 0.99 |
| Fmr1-/y:LS vs. Fmr1-/y:LS+FS | -3 | -9.7 to 3.7 | No | ns | 0.75 |

**Supplementary Table 13. Two-way ANOVA analysis of cFOS cell density quantification in the LaVM subnuclei of the amygdala.**

Summary table of 2-way ANOVA results and Tukey’s multiple comparisons test. cFOS cell density was measured under naïve (no stimulus), exposure to flashing light stimuli (LS), and exposure to LS combined with a footshock (LS+FS) experimental conditions. 2-way ANOVA (alpha 0.05) results show the percentage of variation (%), P-value, P value summary and significant differences at a threshold of $P < 0.05$ per interaction, the sum of squares (SS), mean squares (MS), degrees of freedom (DS), F (DFn, DFd), and P value of the interaction between genotype vs condition, the genotype and the experimental condition effects. Tukey's multiple comparisons post hoc test results are shown for each pairwise comparison, including predicted mean differences, 95% CI of the difference, threshold and summary, and adjusted P-values.

| CeC subnuclei | | | | | |
|---|---|---|---|---|---|
| *Source of Variation* | *% of total variation* | *P value* | *P value summary* | *Significant?* | |
| Interaction | 0.17 | 0.9701 | ns | No | |
| Genotype | 9.7 | 0.0789 | ns | No | |
| Treatment | 23 | 0.0319 | * | Yes | |
| ANOVA table | SS (Type III) | DF | MS | F (DFn, DFd) | P value |
| Interaction | 0.36 | 2 | 0.18 | F (2, 24) = 0.030 | P=0.9701 |
| Genotype | 20 | 1 | 20 | F (1, 24) = 3.4 | P=0.0789 |
| Treatment | 48 | 2 | 24 | F (2, 24) = 4.0 | P=0.0319 |
| Residual | 144 | 24 | 6 | | |
| *Tukey's multiple comparisons test* | *Predicted (LS) mean diff.* | *95.00% CI of diff.* | *Below threshold?* | *Summary* | *Adjusted P Value* |
| WT:Naive vs. WT:LS | 1.2 | -3.4 to 5.8 | No | ns | 0.96 |
| WT:Naive vs. WT:LS+FS | -2 | -6.9 to 2.9 | No | ns | 0.79 |
| WT:Naive vs. Fmr1-/y :Naive | 1.4 | -3.2 to 5.9 | No | ns | 0.94 |
| WT:Naive vs. Fmr1-/y:LS | 3 | -1.6 to 7.5 | No | ns | 0.38 |
| WT:Naive vs. Fmr1-/y:LS+FS | -0.17 | -4.8 to 4.4 | No | ns | >0.9999 |
| WT:LS vs. WT:LS+FS | -3.3 | -8.3 to 1.8 | No | ns | 0.38 |
| WT:LS vs.Fmr1-/y:Naive | 0.14 | -4.7 to 4.9 | No | ns | >0.9999 |
| WT:LS vs. Fmr1-/y:LS | 1.7 | -3.1 to 6.5 | No | ns | 0.87 |
| WT:LS vs. Fmr1-/y:LS+FS | -1.4 | -6.2 to 3.4 | No | ns | 0.94 |
| WT:LS+FS vs. Fmr1-/y:Naive | 3.4 | -1.7 to 8.5 | No | ns | 0.34 |
| WT:LS+FS vs. Fmr1-/y:LS | 5 | -0.090 to 10 | No | ns | 0.06 |
| WT:LS+FS vs.Fmr1-/y:LS+FS | 1.9 | -3.2 to 7.0 | No | ns | 0.86 |
| Fmr1-/y:Naive vs. Fmr1-/y:LS | 1.6 | -3.2 to 6.4 | No | ns | 0.90 |
| Fmr1-/y:Naive vs. Fmr1-/y:LS+FS | -1.5 | -6.3 to 3.3 | No | ns | 0.92 |
| Fmr1-/y:LS vs. Fmr1-/y:LS+FS | -3.1 | -7.9 to 1.7 | No | ns | 0.36 |

**Supplementary Table 14. Two-way ANOVA analysis of cFOS cell density quantification in the CeC subnuclei of the amygdala.**

Summary table of 2-way ANOVA results and Tukey's multiple comparisons test. cFOS cell density was measured under naïve (no stimulus), exposure to flashing light stimuli (LS), and exposure to LS combined with a footshock (LS+FS) experimental conditions. 2-way ANOVA (alpha 0.05) results show the percentage of variation (%), P-value, P value summary and significant differences at a threshold of $P < 0.05$ per interaction, the sum of squares (SS), mean squares (MS), degrees of freedom (DS), F (DFn, DFd), and P value of the interaction between genotype vs condition, the genotype and the experimental condition effects. Tukey's multiple comparisons post hoc test results are shown for each pairwise comparison, including predicted mean differences, 95% CI of the difference, threshold and summary, and adjusted P-values.

| CeM subnuclei | | | | | |
|---|---|---|---|---|---|
| Source of Variation | % of total variation | P value | P value summary | Significant? | |
| Interaction | 5.8 | 0.4263 | ns | No | |
| Genotype | 0.24 | 0.7887 | ns | No | |
| Treatment | 14 | 0.1445 | ns | No | |
| ANOVA table | SS (Type III) | DF | MS | F (DFn, DFd) | P value |
| Interaction | 21 | 2 | 11 | F (2, 24) = 0.88 | P=0.4263 |
| Genotype | 0.88 | 1 | 0.88 | F (1, 24) = 0.073 | P=0.7887 |
| Treatment | 51 | 2 | 25 | F (2, 24) = 2.1 | P=0.1445 |
| Residual | 289 | 24 | 12 | | |
| Tukey's multiple comparisons test | Predicted (LS) mean diff. | 95.00% CI of diff. | Below threshold? | Summary | Adjusted P Value |
| WT:Naive vs. WT:LS | -0.3 | -6.8 to 6.2 | No | ns | >0.9999 |
| WT:Naive vs. WT:LS+FS | -1.4 | -8.4 to 5.5 | No | ns | 0.99 |
| WT:Naive vs. Fmr1-/y :Naive | -0.001 | -6.5 to 6.5 | No | ns | >0.9999 |
| WT:Naive vs. Fmr1-/y:LS | 2.3 | -4.2 to 8.8 | No | ns | 0.88 |
| WT:Naive vs. Fmr1-/y:LS+FS | -3 | -9.5 to 3.5 | No | ns | 0.71 |
| WT:LS vs. WT:LS+FS | -1.1 | -8.3 to 6.1 | No | ns | 1.00 |
| WT:LS vs.Fmr1-/y:Naive | 0.3 | -6.5 to 7.1 | No | ns | >0.9999 |
| WT:LS vs. Fmr1-/y:LS | 2.6 | -4.2 to 9.4 | No | ns | 0.84 |
| WT:LS vs. Fmr1-/y:LS+FS | -2.7 | -9.5 to 4.1 | No | ns | 0.82 |
| WT:LS+FS vs. Fmr1-/y:Naive | 1.4 | -5.8 to 8.6 | No | ns | 0.99 |
| WT:LS+FS vs. Fmr1-/y:LS | 3.7 | -3.5 to 11 | No | ns | 0.60 |
| WT:LS+FS vs.Fmr1-/y:LS+FS | -1.6 | -8.8 to 5.6 | No | ns | 0.98 |
| Fmr1-/y:Naive vs. Fmr1-/y:LS | 2.3 | -4.5 to 9.1 | No | ns | 0.89 |
| Fmr1-/y:Naive vs. Fmr1-/y:LS+FS | -3 | -9.8 to 3.8 | No | ns | 0.75 |
| Fmr1-/y:LS vs. Fmr1-/y:LS+FS | -5.3 | -12 to 1.5 | No | ns | 0.19 |

**Supplementary Table 15. Two-way ANOVA analysis of cFOS cell density quantification in the CeM subnuclei of the amygdala.**

Summary table of 2-way ANOVA results and Tukey's multiple comparisons test. cFOS cell density was measured under naïve (no stimulus), exposure to flashing light stimuli (LS), and exposure to LS combined with a footshock (LS+FS) experimental conditions. 2-way ANOVA (alpha 0.05) results show the percentage of variation (%), P-value, P value summary and significant differences at a threshold of $P < 0.05$ per interaction, the sum of squares (SS), mean squares (MS), degrees of freedom (DS), F (DFn, DFd), and P value of the interaction between genotype vs condition, the genotype and the experimental condition effects. Tukey's multiple comparisons post hoc test results are shown for each pairwise comparison, including predicted mean differences, 95% CI of the difference, threshold and summary, and adjusted P-values.

| *CeL subnuclei* | | | | | |
|---|---|---|---|---|---|
| *Source of Variation* | *% of total variation* | *P value* | *P value summary* | *Significant?* | |
| Interaction | 1.3 | 0.8058 | ns | No | |
| Genotype | 4.2 | 0.2469 | ns | No | |
| Treatment | 25 | 0.0265 | * | Yes | |
| ANOVA table | SS (Type III) | DF | MS | F (DFn, DFd) | P value |
| Interaction | 0.61 | 2 | 0.3 | F (2, 24) = 0.22 | P=0.8058 |
| Genotype | 2 | 1 | 2 | F (1, 24) = 1.4 | P=0.2469 |
| Treatment | 12 | 2 | 5.9 | F (2, 24) = 4.2 | P=0.0265 |
| Residual | 33 | 24 | 1.4 | | |
| *Tukey's multiple comparisons test* | *Predicted (LS) mean diff.* | *95.00% CI of diff.* | *Below threshold?* | *Summary* | *Adjusted P Value* |
| WT:Naive vs. WT:LS | 0.79 | -1.4 to 3.0 | No | ns | 0.87 |
| WT:Naive vs. WT:LS+FS | -1.1 | -3.5 to 1.2 | No | ns | 0.68 |
| WT:Naive vs. Fmr1-/y :Naive | 0.37 | -1.8 to 2.6 | No | ns | 1.00 |
| WT:Naive vs. Fmr1-/y:LS | 1 | -1.2 to 3.3 | No | ns | 0.69 |
| WT:Naive vs. Fmr1-/y:LS+FS | -0.2 | -2.4 to 2.0 | No | ns | 1.00 |
| WT:LS vs. WT:LS+FS | -1.9 | -4.4 to 0.53 | No | ns | 0.19 |
| WT:LS vs.Fmr1-/y:Naive | -0.43 | -2.7 to 1.9 | No | ns | 0.99 |
| WT:LS vs. Fmr1-/y:LS | 0.25 | -2.1 to 2.6 | No | ns | 1.00 |
| WT:LS vs. Fmr1-/y:LS+FS | -0.99 | -3.3 to 1.3 | No | ns | 0.76 |
| WT:LS+FS vs. Fmr1-/y:Naive | 1.5 | -0.95 to 3.9 | No | ns | 0.43 |
| WT:LS+FS vs. Fmr1-/y:LS | 2.2 | -0.27 to 4.6 | No | ns | 0.10 |
| WT:LS+FS vs.Fmr1-/y:LS+FS | 0.93 | -1.5 to 3.4 | No | ns | 0.85 |
| Fmr1-/y:Naive vs. Fmr1-/y:LS | 0.68 | -1.6 to 3.0 | No | ns | 0.94 |
| Fmr1-/y:Naive vs. Fmr1-/y:LS+FS | -0.57 | -2.9 to 1.7 | No | ns | 0.97 |
| Fmr1-/y:LS vs. Fmr1-/y:LS+FS | -1.2 | -3.6 to 1.1 | No | ns | 0.56 |

**Supplementary Table 16. Two-way ANOVA analysis of cFOS cell density quantification in the CeL subnuclei of the amygdala.**

Summary table of 2-way ANOVA results and Tukey's multiple comparisons test. cFOS cell density was measured under naïve (no stimulus), exposure to flashing light stimuli (LS), and exposure to LS combined with a footshock (LS+FS) experimental conditions. 2-way ANOVA (alpha 0.05) results show the percentage of variation (%), P-value, P value summary and significant differences at a threshold of $P < 0.05$ per interaction, the sum of squares (SS), mean squares (MS), degrees of freedom (DS), F (DFn, DFd), and P value of the interaction between genotype vs condition, the genotype and the experimental condition effects. Tukey's multiple comparisons post hoc test results are shown for each pairwise comparison, including predicted mean differences, 95% CI of the difference, threshold and summary, and adjusted P-values.

| Chemical | Catalogue number | Supplier |
| --- | --- | --- |
| TETRAHYDROFURAN, 99%, EXTRA PURE, ANHYDROUS, STABILISED WITH BHT | 10317270 | Thermofisher |
| TETRAHYDROFURAN，REAGENT GRADE 99% | 178810 1L | Merck |
| TETRAHYDROFURAN, 250 PPM BHT | 360589-1L | Merck |
| 2-METHYLTETRAHYDROFURAN BIORENEWABLE | 155810-100ML | Merck |
| 2-METHYLTETRAHYDROFURAN BIORENEWABLE | 414247-1L | Merck |
| TETRAHYDROFURAN ANHYDROUS | 5895700250 | Merck |
| TETRAHYDROFURAN CONTAINS 250 PPM BHT | 87368-1L-M | Merck |
| CYCLOPENTYL METHYL ETHER | 675970-100ML | Merck |
| 1,3-DICHLOROBENZENE | 35350-100ML | Merck |

**Supplementary Table 17. THF and alternative chemicals incompatible with RatDISCO.**
List of THF and other chemicals (name, catalogue number and supplier) tested and that are not compatible with our RatDISCO protocol. They resulted in cloudy tissue unsuitable for light-sheet microscopy.
